# Antibody-drug conjugate combination therapy targeting LGR5 and MET with different payloads enhances efficacy in preclinical colorectal cancer models

**DOI:** 10.64898/2026.07.18.739355

**Authors:** Shraddha Subramanian, Peyton C. High, Cara Guernsey-Biddle, Maya G. Cappellino, Sean P. Sullivan, Zhengdong Liang, Adela M. Aldana, Li Li, Sheng Pan, Kendra S. Carmon

**Author notes:** **Corresponding author:** Kendra S. Carmon University of Texas Health Science Center at Houston The Brown Foundation Institute of Molecular Medicine Center for Translational Cancer Research 1825 Pressler St, SRB 330G Houston, TX 77030. **Authors’ Disclosures:** K.S.C serves on an advisory board for Merus N.V. and Genmab. All other authors declare no potential conflicts of interest.

## Abstract

Leucine-rich repeat-containing G protein-coupled receptor 5 (LGR5) is a marker of cancer stem-like cells frequently upregulated in colorectal cancer (CRC) with lower expression in normal tissues, making it a favorable target for antibody-drug conjugates (ADCs). ADCs combine antibody specificity with potent cytotoxic payloads to enhance efficacy while minimizing systemic toxicity. LGR5-targeting ADCs incorporating different payloads demonstrate strong initial tumor inhibition, yet tumors eventually recur due in part to LGR5 downregulation, necessitating more effective strategies to prevent relapse. We show treatment with chemotherapies or an LGR5-targeting ADC coupled to a topoisomerase 1 inhibitor payload (8E11-CPT2) reduces LGR5 levels and increases MET receptor expression and/or activation in CRC cells, supporting a therapeutic approach to target LGR5 and MET simultaneously. Accordingly, we engineered a MET-targeting ADC (ABT-700-SG3199) via site-specific conjugation of the anti-MET antibody telisotuzumab (ABT-700) with the DNA-crosslinking pyrrolobenzodiazepine (PBD) dimer SG3199. ABT-700-SG3199 exhibited superior potency and efficacy in CRC models compared to the clinical-stage MET-targeting ADCs ABBV-399 and ABBV-400, which use the same antibody backbone conjugated to different payloads. Treatment with ABT-700-SG3199 increased LGR5 expression, and the combination of ABT-700-SG3199 with 8E11-CPT2 enhanced CRC cell-killing efficacy, reinforcing the rationale for a dual-targeting approach. In patient-derived xenografts, combined administration of 8E11-CPT2 and ABT-700-SG3199 markedly delayed tumor relapse and prolonged survival compared with single-agent treatment. Taken together, these findings reveal a reciprocal regulation between MET and LGR5 in response to LGR5- or MET-targeting ADCs in CRC models and support a LGR5/MET dual-targeting therapeutic strategy to enhance efficacy and potentially overcome resistance and relapse.

## Introduction

Antibody-drug conjugates (ADCs) are a rapidly expanding class of targeted therapeutics revolutionizing the clinical landscape of precision oncology. ADCs act as biological “Trojan Horses,” by attaching a monoclonal antibody (mAb) that specifically targets antigens overexpressed on the tumor cell surface to a linker-payload moiety, enabling the efficient delivery of potent cytotoxins into cancer cells while sparing healthy tissues (1). This modular design confers numerous advantages over conventional chemotherapy, including increased tumor specificity, reduced systemic toxicity, and enhanced efficacy (1). Though several ADCs have been FDA-approved for the treatment of various solid and hematological malignancies, none have been specifically approved for colorectal cancer (CRC), which remains a leading cause of cancer-related mortality largely due to resistance-driven metastatic relapse (2). The HER2-targeting ADC, trastuzumab deruxtecan (T-Dxd), has received tumor-agnostic approval for HER2-expressing solid tumors (3,4). However, the clinical benefit in CRC is limited to HER2-amplified tumors, which account for only 3-5% of patients (5). These limitations highlight the need for better target antigen selection for the development of ADCs to improve treatment outcomes in CRC.

Leucine-rich repeat-containing G protein–coupled receptor 5 (LGR5) is a bona fide marker of cancer stem-like cells (CSCs), a subpopulation of cells defined by heightened tumorigenic potential and intrinsic drug resistance (6,7). LGR5^⁺^ CSCs dynamically shift between proliferative LGR5^⁺^ and undifferentiated LGR5^⁻^ states to evade therapeutic stresses and drive metastasis (8–10). We and others developed LGR5-targeting ADCs conjugated to monomethyl auristatin E (8F2-MMAE) that induced marked tumor regression in some CRC models (11,12). However, relapse occurred after treatment withdrawal, due in part to resistance mechanisms driven by CRC cell plasticity, target downregulation, and suboptimal payload (10,12). Our more recent LGR5-directed ADC (8E11-CPT2), incorporating a topoisomerase 1 inhibitor (TOP1i) payload (camptothecin derivative, CPT2), showed significant potency and efficacy across a broader set of CRC models (13). However, at doses tested, 8E11- CPT2 only transiently delayed progression and was unable to prevent tumor regrowth as a monotherapy (13). These findings reinforce the need for rational combinatorial strategies that target both LGR5^+^ and LGR5^-^ populations to overcome ADC resistance and extend therapeutic durability.

Resistance to antibody-based therapies, including ADCs, is multifactorial, with target downregulation and receptor tyrosine kinase (RTK) activation as potential bypass mechanisms (10,14,15). Activation and/or amplification of MET, an RTK aberrantly expressed in CRC and a known driver of invasion, progression, and metastasis (16,17), has been shown to drive resistance to clinically approved targeted therapies (18). Further, we previously showed that LGR5^+^ CRC cells could evade 8F2-MMAE treatment through LGR5 downregulation and concomitant MET activation, demonstrating a potential mechanism of LGR5-targeting ADC resistance (10). Furthermore, this suggests that co-targeting MET may be a rational strategy to overcome treatment resistance. Although no MET-targeting inhibitors or ADCs have received clinical approval for CRC, telisotuzumab vedotin (ABBV-399), a MET-directed ADC with an MMAE payload, has been approved for non-small cell lung cancer (NSCLC) (19,20). Moreover, telisotuzumab adizutecan (ABBV-400), a MET-targeting ADC with a TOP1i payload, has shown clinical benefit in NSCLC and other cancers, including CRC (21,22).

In this study, we demonstrate that LGR5 downregulation and MET upregulation and/or MET activation occur in response to LGR5-targeting ADCs and standard-of-care chemotherapies. To overcome this potential resistance mechanism, we developed a MET-targeting ADC (ABT-700-SG3199) by site-specific conjugation of the DNA crosslinking agent pyrrolobenzodiazepine (PBD) dimer SG3199 to the MET-targeting mAb ABT-700 (telisotuzumab) (23). ABT-700-SG3199 exhibited high target selectivity and superior potency and efficacy in vitro compared to ABBV-399 and ABBV-400. Further, ABT-700-SG3199 also induced significant tumor growth inhibition in CRC xenografts. However, as with LGR5-directed ADCs, single-agent activity did not prevent relapse. In fact, treatment with MET-targeting therapies increased LGR5 levels, underscoring the rationale for a LGR5/MET dual-targeting approach. Combination treatment with 8E11-CPT2 and ABT-700-SG3199 markedly suppressed tumor regrowth and prolonged survival compared to monotherapy. Collectively, these findings identify a reciprocal regulation of MET and LGR5 as a potential resistance mechanism to LGR5- or MET-targeting ADCs in CRC, respectively, and suggests LGR5/MET dual-targeting ADC therapy may be a promising strategy to overcome resistance and relapse.

## Methods and Materials

### Cell lines

Cell lines procured from ATCC include: DLD-1 (Cat# CCL-221, RRID: CVCL_0248), HCT116 (Cat# CCL-247, RRID: CVCL_0291), HEK293T (Cat# CRL-3216, RRID:CVCL_0063), HT-29 (Cat# HTB-38, RRID: CVCL_0320), LoVo (Cat# CCL- 229, RRID: CVCL_0399), LS180 (Cat# CL-187, RRID: CVCL_0397), and SW620 (Cat# CCL-227, RRID: CVCL_0547). LIM1215 (Cat# 10092301, RRID: CVCL_2574) was obtained from Millipore Sigma. All cell lines were purchased new and/or authenticated by short tandem repeat profiling and confirmed to be free of mycoplasma contamination. HEK293T and CRC cells were cultured in DMEM and RPMI-1640 medium (Gibco, Cat# 10566016 and 11875093), respectively, and supplemented with 10% fetal bovine serum (Gibco, Cat# 26140079) and 1% penicillin-streptomycin (Gibco, Cat# 15140122). LIM1215 cells were supplemented with 10 μM 1-thioglycerol (Sigma, Cat# 8864025), 25 mM HEPES (Gibco, Cat# 15630080), 0.6 μg/mL insulin (Gibco, Cat# 12585014), and 1 μg/mL hydrocortisone (Sigma, Cat# H0888). All cell lines were maintained at 37 °C in a humidified incubator with 5% CO_2_.

### RNAi-based knockdown

For stable shRNA knockdown (KD), DLD-1 and LoVo cells were transduced with lentivirus particles produced by co-transfecting HEK293T cells with a pLKO.1 vector with MET-targeting short hairpin RNA (shRNA; #TRCN0000121090 GCATGTCAACATCGCTCTAAT, Dharmacon) and packaging plasmids, psPAX2 and pMD2.G, using FuGENE**^®^** HD (Promega, Cat# E2311) (10,12). For transient MET silencing, DLD-1 cells were seeded at 40–50% confluency in 6-well plates (GenClone, Cat# 25-105) and transfected with either MISSION^®^ negative control small interfering RNA (siRNA; Sigma, Cat# SIC001) or MET-specific siRNA (FlexiTube siRNA Qiagen, Cat# SI00300874) at a final concentration of 100 nM using JetPRIME (Polyplus, Cat# 55-132).

### Antibody cloning, production, and purification

ABT-700 (telisotuzumab) was produced using publicly available sequences. Variable heavy (V_H_) and light (V_L_) chain regions were subcloned into pCEP4 expression vectors containing human IgG1 or kappa constant domains, respectively, using In- Fusion Snap Assembly (Takara, Cat# 638947). To enable facile site-specific conjugation to Q295, an N297A mutation was introduced into the Fc region to eliminate the need for deglycosylation (24). mAbs were transiently expressed in Expi293F cells (ThermoFisher, Cat# A14527, RRID: CVCL_D615) using polyethylenimine HCL Max (Polysciences, Cat# 247651). Medium was harvested 7 days post-transfection, and mAbs were purified by Protein A affinity chromatography (Genscript, Cat# L00210), eluted with pH 3 glycine, neutralized with pH 8.5 Tris-HCl, and buffer exchanged into PBS. SDS-PAGE and Coomassie staining were performed to confirm mAb purity. Non-targeting hIgG1 isotype control rituximab (cmAb) was generated similarly as previously described (24). Rituximab was used as it targets CD20 on human B-cells and doesn’t bind mouse antigen. MAbs were quantified using a NanoDrop (Thermo Fisher).

### ADC production

ABT-700 and cmAb were conjugated at the Q295 site using an enzymatic transamidation strategy, as we previously reported (13,24). MAbs were incubated at ambient temperature with 40 molar equivalents of an amino-PEG4-azide linker (BroadPharm, Cat# BP-21615) and Activa^®^ TI microbial transglutaminase (mTG; 495 mg; 8% final concentration; Modernist Pantry) for 8 hours to facilitate site-specific incorporation of the azide moiety. Post-reaction, unreacted linker and residual enzyme were removed by Protein A affinity chromatography. The linker-payload DBCO-PEG8-VA-PAB-SG3199 (Levena Biopharma, Cat# SET0317) was prepared at 10 mg/mL in DMSO and added at a stoichiometric ratio of 1.5 equivalents per azide. The strain-promoted azide–alkyne cycloaddition reaction was performed at ambient temperature for 4 hours to enable site-specific conjugation of the linker-payload to each azide. The reaction mixture was buffer exchanged into 5 mmol/L phosphate buffer (pH 6.5) and purified by Protein A affinity chromatography. ABT- 700-SG3199 and cADC were eluted and buffer-exchanged into PBS and recovered at concentrations of 3.84 mg/mL (29.25 mg total; 75% yield) and 4.61 mg/mL (17.46 mg total; 68% yield), respectively. 8F2-SG3199 ADC comprised of anti-LGR5 (8F2) mAb conjugated to PEG8-VA-PAB-SG3199, 8E11-SG3199 ADC composed of anti- LGR5 (8E11) mAb linked to PEG8-VA-PAB-SG3199, and 8E11-CPT2 ADC consisting of 8E11 mAb attached to DBCO-PEG8-VKG-CPT2 were generated with DARs = 2.0, 2.0 and 7.94, respectively, as previously reported (13,25).

### Mass spectrometry

Drug-to-antibody ratio (DAR) determination of ADCs was performed at the UTHealth Houston Clinical and Translational Proteomics Center using an Agilent 6538 UHD Accurate-Mass Q-TOF LC/MS system coupled to an Agilent 1200 HPLC. Samples were diluted in a 25% acetonitrile, 75% water solution with 0.1% formic acid before injection to optimize ionization efficiency. Chromatographic separation was achieved on an Agilent PLRP-S reversed-phase column (2.1 × 50 mm, 5 μm, 1000 Å) under a linear gradient of 25–90% solvent B (80% acetonitrile, 20% water, 0.1% formic acid) over 20 minutes at a flow rate of 0.2 mL/min. Mass spectrometric detection was performed in electrospray ionization (ESI) positive mode, with a capillary voltage of 3500 V, a drying gas flow of 7 L/min, and a source temperature of 325 °C, scanning across an m/z range of 650–2800. Data acquisition was performed in MS1 scan mode between 4.5 and 20 minutes. Raw spectra were processed using Agilent MassHunter BioConfirm (v. B. 04.00), employing Maximum Entropy deconvolution to generate intact mass profiles within a 20-100 kDa range, enabling the determination of average DAR = 1.6 for ABT-700-SG3199 and DAR = 1.5 for cADC.

### Internalization assays

CRC cells were seeded onto poly-D-lysine–coated 8-well chamber slides (Corning, Cat# 354632) and allowed to adhere overnight. For lysosomal co-localization studies, cells were incubated with ABT-700 (10 μg/mL) at 37 °C for 1 hour, followed by fixation in 4% methanol-free formaldehyde (Thermo Fisher, Cat# 28906) and permeabilization with 0.1% saponin (Sigma, Cat# 84510). Cells were subsequently stained with anti-LAMP1 (1:400; Cell Signaling Technology (CST), Cat# 9091, RRID: AB_2687579) and detected using Alexa Fluor 488-conjugated goat anti-rabbit IgG (1:200; ThermoFisher, Cat# A-32731, RRID: AB_2633280) along with Alexa Fluor 555-conjugated goat anti-human IgG (1:200; ThermoFisher, Cat# A-21433, RRID: AB_2535854). For LGR5/MET co-localization, cells were incubated for 2 hours with rat anti-LGR5 (clone 8F2, 5 μg/mL; BD, Cat# 562731) and human anti-MET (ABT- 700, 10 μg/mL) mAbs, followed by fixation, permeabilization, and incubation with Alexa Fluor 488–conjugated rabbit anti-rat IgG (1:200; ThermoFisher, Cat# A-21210, RRID: AB_2535796) and Alexa Fluor 555–conjugated goat anti-human IgG. Nuclear counterstaining was performed using TO-PRO-3 (1:1000; Thermo Fisher, Cat# T3605). Images were acquired using a Leica TCS SP5 confocal microscope (Leica Microsystems, RRID: SCR_020233) and processed with LAS AF Lite software.

### Fluorescence-based cell binding assays

CRC cells were seeded into black poly-D-lysine–coated 96-well plates (Corning, Cat# 354407) and allowed to adhere overnight or until confluent. To detect surface binding, serial dilutions of cmAb, ABT-700, cADC, and ABT-700-SG3199 were added to cells and incubated at 4 °C for 2 hours. Plates were washed with PBS, fixed with 4% paraformaldehyde, and incubated with Alexa Fluor 555-conjugated goat anti-human IgG (1:400) for 1 hour at room temperature. After a series of PBS washes, fluorescence intensity was quantified using an Infinite M1000 plate reader (Tecan, RRID: SCR_025732). Reported K_d_ values are the average of 2-3 independent experiments in triplicate.

### Cell viability assays

CRC cells were seeded at a density of 1,000 cells per well in 96-half-well white plates (Corning, Cat# 3688) and allowed to adhere overnight. Cells were treated with serial dilutions of cmAb, ABT-700, cADC, ABT-700-SG3199, Telisotuzumab vedotin (ABBV-399; MedChemExpress, HY-141601), Telisotuzumab adizutecan (ABBV-400; MedChemExpress, HY-171945), 8E11-CPT2 (13), or 8F2-SG3199 (25) as single agents or in combination. After 5 days of incubation at 37°C, cell viability was quantified using the CellTiter-Glo^®^ 2.0 assay (Promega, Cat# G9242). Luminescence was measured using the Infinite M1000 plate reader (Tecan, RRID: SCR_025732). IC_50_ values represent the mean of 2-3 independent experiments in triplicate. Drug interactions were analyzed using the Loewe Additivity Model implemented in SynergyFinder+ (26). Synergy plots represent the mean of three independent experiments, with cell viability for select drug combinations plotted as individual quadruplicate data points from each.

### Western Blot

Whole-cell lysates were prepared using RIPA buffer (Thermo Fisher, Cat# 89901) supplemented with Halt™ protease and phosphatase inhibitors (Thermo Fisher, Cat# 78443). For patient-derived xenograft (PDX) tissues, samples were homogenized, subjected to freeze–thaw, sonicated, and centrifuged at 13,000 rpm for 10 min at 4 °C. Protein concentrations were quantified using the BCA assay (ThermoFisher, Cat# 23227). Lysates were denatured in SDS Laemmli buffer with β- mercaptonethanol at 37 °C for 1 hour before SDS-PAGE. Antibodies used include: anti-mouse IgG (1:2000; CST, Cat# 7076, RRID: AB_330924), anti-rabbit IgG (1:2000; CST, Cat# 7074, RRID: AB_2099233), anti-MET (1:1000; CST, Cat# 8198, RRID: AB_10858224), anti-p-MET Y1234/1235 (1:1000; CST, Cat# 3077, RRID: AB_2143884), anti-LGR5 (1:1000; Abcam, Cat# ab75732, RRID: AB_1310281), and anti–β-actin (1:10,000; CST, Cat# 3700, RRID: AB_2242334). Chemical inhibitors used include: 5-fluorouracil (5-FU; Acros, Cat# 10722125), SN-38 (Tocris, Cat# 2684), and crizotinib (CST, Cat# 4401).

### In Vivo Efficacy Studies

All animal experiments were approved by the UTHealth Houston Institute of Molecular Medicine (IMM) IACUC (AWC-23-0106). The patient-derived xenograft (PDX) model studies were performed in compliance with ethical guidelines (i.e., Declaration of Helsinki, Belmont Report, U.S. Common Rule), and appropriate approvals were obtained by the UTHealth Houston Institutional Review Board (HSC-MS-20-0327, HSC-MS-21-0074). PDX tumor fragments (2-3 mm) from CRC-001 or XST-GI-010 models (13,24) were implanted in NOD/SCID gamma (NSG) mice (Jackson Laboratory, RRID: IMSR_JAX:005557). CRC cell line-derived xenograft (CDX) models were established by subcutaneous injection of 1 × 10 HCT116 cells or 2 × 10 DLD1 cells in nu/nu mice (Charles River, RRID: IMSR_CRL:088; Jackson Laboratory, RRID: IMSR_JAX:002019). For all studies, 6- to 8-week-old female mice were used. Once tumors reached ∼100-150 mm³, mice were randomized and treated with a single dose or dosed once weekly for 2 weeks intraperitoneally (IP) with vehicle (PBS), cmAb, cADC, ABT-700-SG3199, 8E11-CPT2, or ABT-700-SG3199 combined with 8E11-CPT2. Tumor volumes were measured biweekly using calipers and calculated as (length × width²) / 2. Mice were euthanized when tumors reached 15 mm in diameter.

### Toxicity Study

Female C57BL/6 mice (6-8 weeks old; *n* = 3-4 per group; Charles River, RRID:IMSR_CRL:027) were administered a single IP dose of ABT-700-SG3199 at 0.1, 0.5, or 1.5 mg/kg, or PBS as vehicle control. Animals were monitored for body weight every 2-3 days for 2 weeks and predefined humane endpoints (i.e., >20% weight loss or signs of severe distress) were not met. At study termination, mice were anesthetized, and whole blood was collected via cardiac puncture for hematology and clinical chemistry analyses in the Veterinary Medicine & Surgery Department at MD Anderson Cancer Center. Liver and kidney tissues were harvested, fixed, paraffin-embedded, sectioned, and stained with hematoxylin and eosin (H&E) by the UTHealth Houston IMM histopathology core for histopathological evaluation.

### Statistical analysis

All data were analyzed using GraphPad Prism (RRID: SCR_002798) and are presented as mean ± SD unless otherwise specified. In vitro binding and viability curves were fitted using a nonlinear logistic regression model. For transcriptomic comparisons, matched tumor and normal samples from The Cancer Genome Atlas (TCGA) were analyzed using paired Student’s *t*-tests. Statistical significance of in vitro assays was assessed using one-way ANOVA followed by Tukey’s multiple-comparison test. Efficacy studies were analyzed using one-way ANOVA with Tukey’s post hoc test for multiple group comparisons or Student’s *t*-test for pairwise comparisons. Survival analyses were performed using Kaplan–Meier plots with log-rank tests to evaluate treatment effect on overall survival. A *p-value* < 0.05 was considered statistically significant.

## Results

### LGR5 and MET are highly expressed in primary and metastatic colorectal tumors

Prior to investigating the interplay between LGR5 and MET in response to standard-of-care and LGR5-targeting therapies, we performed a side-by-side analysis of gene expression levels in CRC patient datasets and cell lines. RNA-sequencing (RNA-seq) data from The Cancer Genome Atlas (TCGA) colorectal adenocarcinoma (COADREAD) cohort showed significant upregulation of LGR5 and MET **(Fig. 1A)** in tumors compared with matched adjacent normal epithelium (n=32, *p*<0.0001), consistent with other reports (11,12,16). Similarly, an independent dataset of 18 paired primary tumors, synchronous liver metastases, and normal colonic mucosa revealed elevated expression of both genes in primary and metastatic lesions **(Figs. 1B-C)**(27). Western blot analysis and quantification confirmed high LGR5 and/or MET expression in a panel of CRC cell lines consistent with RNA-seq data from the Cancer Cell Line Encyclopedia (CCLE) (**Figs. 1D-F**).

**Figure 1.**
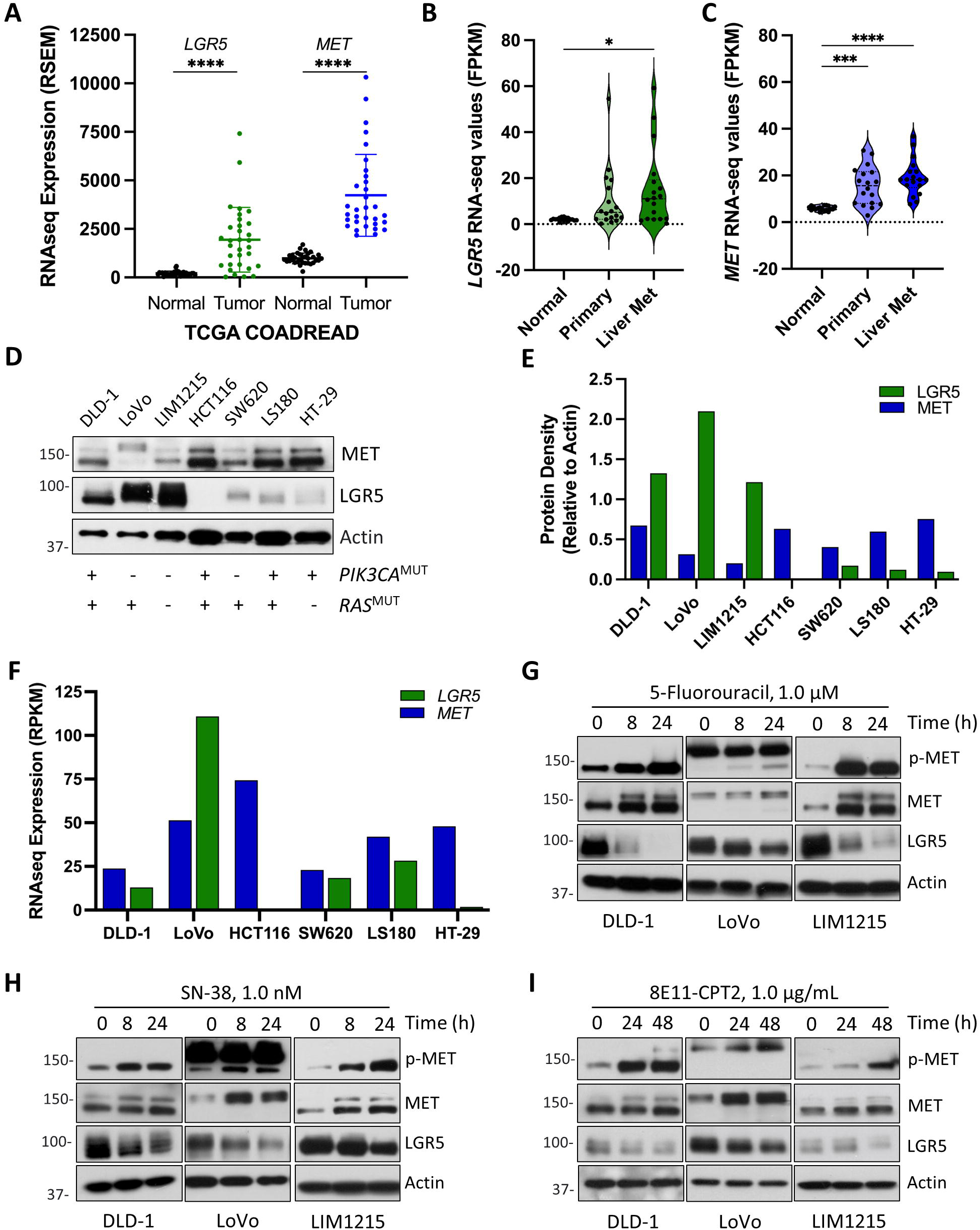
LGR5 and MET are highly expressed in CRC and are differentially regulated following chemotherapy or LGR5-targeted ADC treatment. **(A)** LGR5 and MET RNA-seq expression in normal intestinal mucosa and adjacent colorectal tumors from the TCGA COADREAD cohort (*n* = 32). Values represent RNA-Seq by Expectation-Maximization (RSEM). RNA-seq expression of **(B)** LGR5 and **(C)** MET in an independent cohort of 18 matched samples comprising normal colonic epithelium, primary colorectal adenocarcinoma, and synchronous liver metastases (GSE50760). Values represent Fragments Per Kilobase of transcript per Million (FPKM). **(D)** Western blot and **(E)** quantification of endogenous MET and LGR5 protein expression across a panel of CRC cell lines. **(F)** RNA-seq expression of LGR5 and MET from the Cancer Cell Line Encyclopedia (CCLE). Values represent reads per kilobase per million (RPKM). **(G-I)** Western blots of total MET, phospho-MET and LGR5 expression in DLD-1, LoVo, and LIM1215 cells following treatment with **(G)** 5-Fluorouracil (5-FU, 1 µM), **(H)** SN-38 (1 nM), or (I) LGR5 ADC (8E11-CPT2, 1 µg/mL) over the indicated time course. Results represent ≥3 independent experiments. Statistical significance determined using paired student’s t-test or ANOVA (*p* *<0.05, *p* ***<0.001, and *p* ****< 0.0001). Quantitative data are presented as mean ± SD.

### Chemotherapies and LGR5-targeting ADCs decrease LGR5 levels and increase MET activation in CRC cells

Next, we evaluated changes in LGR5 and MET levels in response to chemotherapies in CRC cell lines of various genetic backgrounds. LoVo (KRAS^MUT^/PIK3CA^WT^), DLD-1 (KRAS^MUT^/PIK3CA^MUT^), and LIM1215 (KRAS^WT^/PIK3CA^WT^) CRC cells were treated with SN-38 (the active metabolite of irinotecan) or 5-FU at concentrations below IC_25_ (**Supplementary Figs. S1A-B)**. Western blot analysis revealed a time-dependent reduction in LGR5 expression, accompanied by increases in total and phosphorylated MET (**Fig. 1G-H**). Noticeably, both the partially glycosylated precursor (∼170 kDa) and mature β-subunit (∼145 kDa) of MET were increased in DLD-1 and LIM1215 cells (**Figs. 1G-H**). LoVo cells showed an increase in total levels of the aberrantly processed single-chain pro-MET (∼190 kDa) and induction of a smaller (∼145 kDa) processed form with increased phosphorylation over time (**Figs. 1G-H**), consistent with previous results following irinotecan treatment (10). Next, we tested the effects of 8E11-CPT2, our LGR5- targeting ADC incorporating the TOP1i payload CPT2 (DAR=8) (13). 8E11-CPT2 exhibited significant efficacy in CRC models with IC_50_ values in the low nanomolar range (13). Like chemotherapies, 8E11-CPT2 treatment decreased LGR5 expression and increased total MET and its phosphorylation after 48 hours (**Fig. 1I**). These data show loss of chemotherapy- and LGR5-targeting ADC-induced loss of LGR5 expression corresponds with MET upregulation and/or activation independent of *KRAS* or *PI3KCA* mutations. Moreover, these results suggest targeting MET in combination with these therapies may be a more effective approach to treat CRC.

### Generation and characterization of a MET-targeting ADC, incorporating DNA-damaging PBD payload

To improve upon existing MET-targeting ADC strategies, we generated a MET-targeting ADC incorporating a valine–alanine (VA) dipeptide cleavable linker and equipped with the more potent payload SG3199 (**Fig. 2A**). SG3199 is a PBD that induces covalent DNA interstrand crosslinks and has shown to exhibit sub-nanomolar potency across cancer cell lines (28). We produced, purified, and verified ABT-700 **(Supplementary Figs. S2A-B)**, a clinical-stage, bivalent mAb that binds and disrupts dimerization of human (and not murine) MET and blocks both hepatocyte growth factor -dependent and -independent signaling (29). A non-targeting isotype control mAb (cmAb) was also generated to detect off-target binding **(Supplementary Fig. S2C)**. ABT-700 was then tested and compared to cmAb for specific binding to MET-expressing CRC cells and effective lysosomal trafficking, which is critical for ADC payload release. Using a fluorescence-based binding assay, we showed ABT-700 binds MET at the surface of DLD-1 cells with high affinity (K_d_ ≈ 0.036 ± 0.010 μg/mL or 0.249 ± 0.087 nM), similar to values previously reported for other cancer cell lines (29). In contrast, cmAb exhibited negligible binding (**Fig. 2B**). Immunocytochemistry (ICC) showed that ABT-700 internalizes and co-localizes with the lysosome marker LAMP1 after 1 hour in HCT116, DLD-1, and LoVo cells (**Fig. 2C and Supplementary Fig. S3A**). No binding was detected for cmAb in HCT116 and LoVo cells. To generate the MET-targeting ADC, site-specific conjugation of ABT-700 was performed using microbial transglutaminase (mTG)-mediated and strain-promoted azide–alkyne cycloaddition as previously reported (13,24), enabling precise attachment of a DBCO-functionalized PEG8-VA-SG3199 linker–payload to Q295 residues in the Fc region (**Fig. 2A**). A non-targeting isotype control ADC (cADC) was generated using the same approach with cmAb. Conjugation of ABT- 700-SG3199 (DAR∼1.6) or cADC (DAR∼1.5) was verified by SDS-PAGE with Coomassie staining and LC-MS (**Supplementary Figs. S2A-C**). Further, ABT-700- SG3199 was retained binding affinity similar to ABT-700 on DLD-1 cells (K_d_ ≈ 0.047 ± 0.018 μg/mL or 0.315 ± 0.121 nM; **Fig. 2B**).

**Figure 2.**
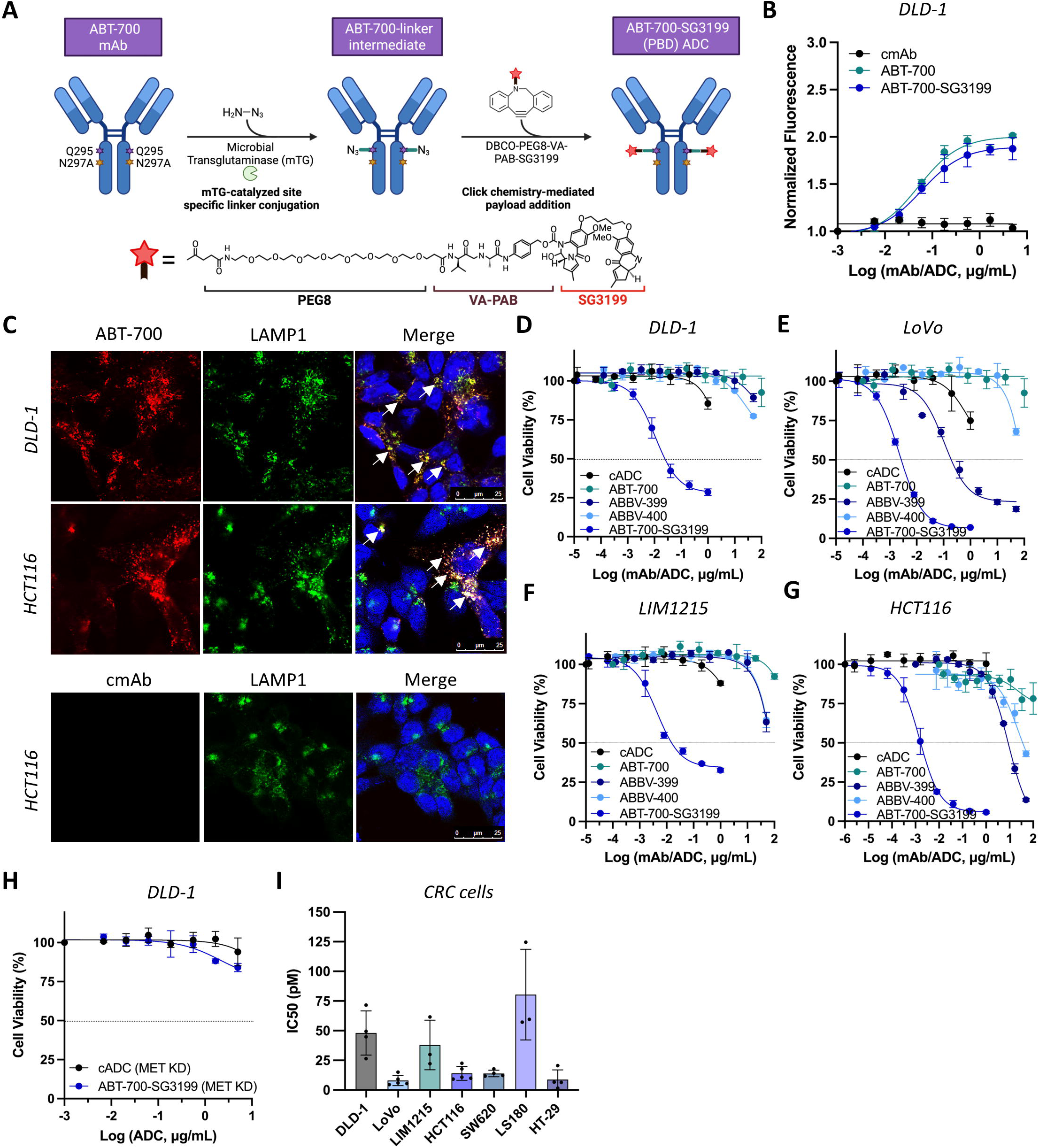
Generation of ABT-700–SG3199 and evaluation of its potency relative to clinical MET ADCs in CRC cells. **(A)** Schematic of microbial transglutaminase (mTG)–mediated conjugation of a cleavable valine-alanine (VA) linker to the Q295 residue in the ABT-700 Fc region, followed by strain-promoted alkyne-azide cycloaddition to attach the PBD dimer payload (SG3199), generating ABT-700- SG3199 (DAR = 1.6). **(B)** DLD-1 cell-based binding assay demonstrating equivalent MET binding by ABT-700 mAb and ABT-700-SG3199. **(C)** Immunocytochemistry showing ABT-700 binding to MET and colocalization (yellow) with the lysosomal marker LAMP1 after 1 hour at 37 °C in DLD-1 and HCT116 cells. cmAb shows no binding. Scale bars: 25 µm. **(D-G)** Dose-dependent cytotoxicity of ABT-700-SG3199 compared with cADC, ABT-700 mAb, ABBV-399 (MMAE) and ABBV-400 (Adizutecan) in **(D)** DLD-1, **(E)** LoVo, **(F)** LIM1215 and **(G)** HCT116 cells, measured 5 days post-treatment. **(H)** Potency of ABT-700-SG3199 and cADC in DLD-1 cells with siRNA-mediated MET knockdown, measured 4 days post-treatment. **(I)** IC_50_ values of ABT-700-SG3199 across the CRC cell line panel. Results represent 2-3 independent experiments. Quantitative data are presented as mean ± SD.

### ABT-700-SG3199 demonstrates increased efficacy compared to clinical-stage and approved MET-targeting ADCs in CRC cells

Next, we evaluated the efficacy of ABT-700-SG3199 compared with cADC, unconjugated ABT-700 mAb, and MET-targeting ADCs ABBV-399 and ABBV-400 in CRC cells expressing different levels and forms of MET. ABBV-399 and ABBV-400 are comprised of ABT-700 conjugated to either MMAE (DAR∼3.1) or a proprietary adizutecan TOP1i (DAR∼6) payload, respectively (19,21). As shown in **Figs. 2D-G**, ABT-700-SG3199 outperformed all other ADCs in CRC cells tested. ABT-700 and ABBV-400 exhibited minimal effects in DLD-1, LoVo, and LIM1215 cells. ABBV-399 demonstrated relatively potent efficacy in LoVo cells (IC_50_ = 0.098 ± 0.010 μg/mL or 0.655 ± 0.067 nM) but exhibited limited activity in DLD-1 and LIM1215 cells. In HCT116 cells, the IC_50_ values for ABBV-399 and ABBV-400 were 7.946 ± 1.197 μg/mL (53.14 ± 8.01 nM) and 20.63 ± 3.71 μg/mL (137.94 ± 24.83 nM), respectively. This was consistent with previous findings showing that LoVo cells were sensitive to MMAE-based ADCs, whereas most CRCs are relatively more resistant (12,30). On the other hand, ABT-700-SG3199 exhibited much higher potency in DLD-1, LoVo, LIM1215, and HCT116 cells **(Figs. 2D-G**). To verify target specificity, we showed that ABT-700-SG3199 had minimal activity in DLD-1 MET KD cells (**Fig. 2H**), confirming MET-dependent cell-killing. MET KD using siRNA was confirmed by western blot (**Supplementary Fig. S3B**). ABT-700-SG3199 was also evaluated in MET-expressing SW620, HT-29, and LS180 cells **Fig. 1D and Supplementary Figs. S3C-E**). Average IC_50_ values for ABT-700-SG3199 ADC in all CRC cell lines tested were in picomolar range shown in **Fig. 2I** and summarized in **Table S1**. The cADC exhibited negligible cytotoxicity in all CRC cell lines tested (**Figs. 2D-H and Supplementary Figs. S3C-E**). The potencies of the different MET ADCs were relatively consistent with the intrinsic sensitivity of each CRC cell line to free payloads, as all lines showed greatest sensitivity to SG3199 (**Supplementary Fig. S3F-J**). Though MMAE and adizutecan showed similar IC_50_ values in LoVo and HCT116 cells, ABBV-399 demonstrated greater potency compared to ABBV-400 despite lower DAR. This suggesting that, in addition to payload potency, the distinct peptide linkers and/or conjugation strategies may also be contributing to differences in ADC efficacy. Overall, our findings show ABT-700-SG3199 has increased potency and efficacy in CRC cells compared to clinically validated MET-targeting ADCs ABBV-399 and ABBV-400.

### ABT-700-SG3199 is well-tolerated and exhibits robust anti-tumor activity in preclinical CRC models

Before evaluating ABT-700-SG3199 efficacy in vivo, we performed a single-dose safety study to assess potential payload-mediated, off-target toxicities in immunocompetent C57BL/6 mice. As ABT-700-SG3199 only binds human MET, on-target off-tumor effects were not examined. Animals received escalating doses of ABT-700-SG3199 (0, 0.1, 0.5, and 1.5 mg/kg), and after 2 weeks, blood was collected for hematology and clinical chemistry analyses and tissues were excised for histopathology. Dosing concentrations were selected based on the payload’s potency and the clinical dosing of PBD-based ADCs (31). Histopathological evaluation by H&E staining revealed no pathological changes in liver or kidney tissue architecture (**Supplementary Figs. S4A-B**) compared to vehicle. Furthermore, no significant changes in body weight (**Fig. 3A**), liver enzymes (aspartate aminotransferase and alanine aminotransferase; **Figs. 3B-C**), kidney function (creatinine; **Fig. 3D**), or white blood cell counts (**Fig. 3E**) were observed at doses tested. Collectively, these findings indicate that ABT-700-SG3199 is well tolerated in mice, with no detectable off-target toxicities.

**Figure 3.**
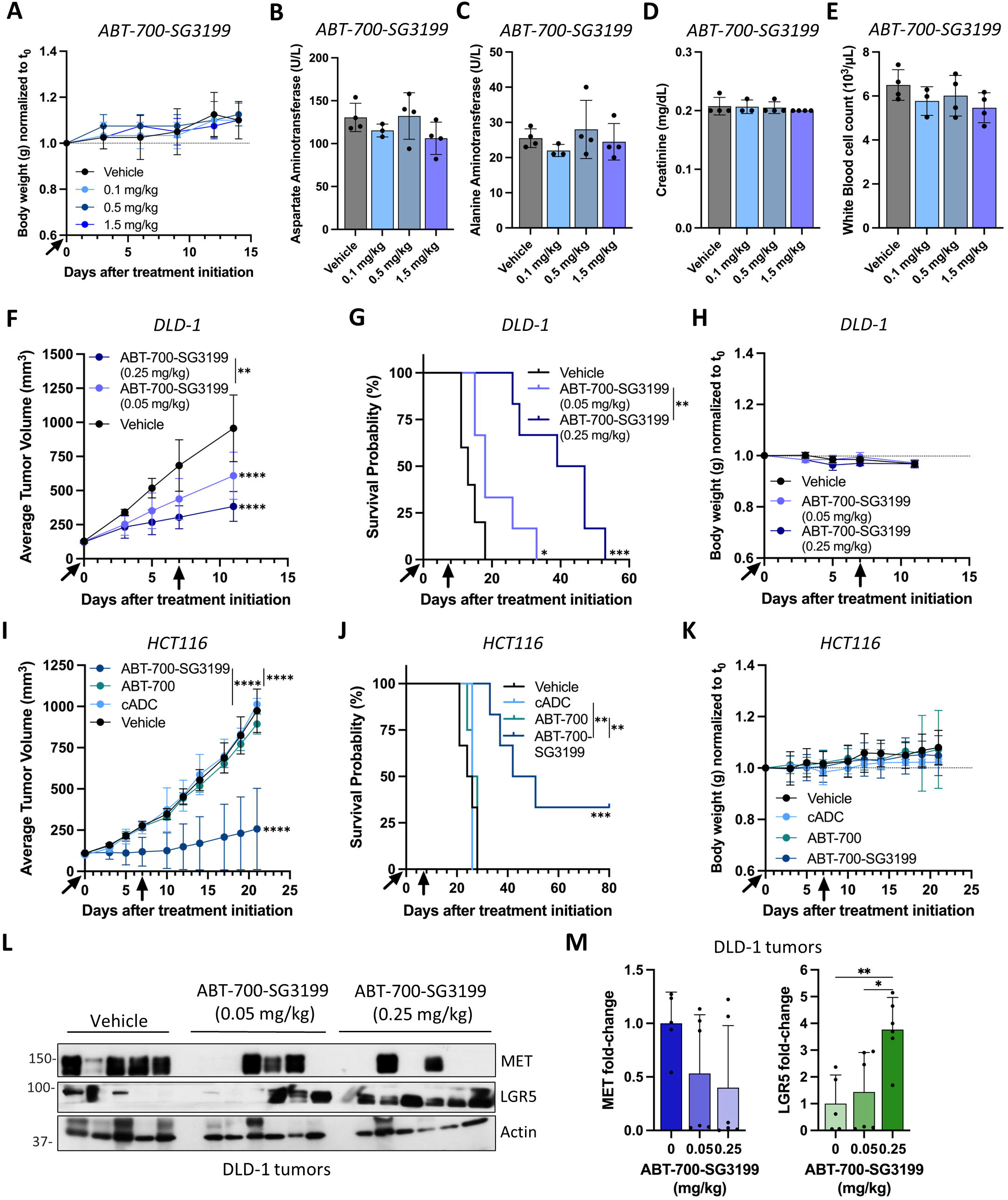
Tolerability and anti-tumor efficacy of ABT-700-SG3199 in CRC cell line-derived xenograft models. **(A)** Body weight changes in C57BL/6 mice (*n* = 3 for the 0.1 mg/kg group; *n* = 4 for all other groups), **(B-C)** serum liver enzymes (aspartate and alanine aminotransferases), **(D)** kidney function marker (creatinine), and **(E)** white blood cell counts 2 weeks post ABT-700-SG3199 treatment. **(F)** DLD-1 CDX tumor growth until day 11 when first vehicle-treated mouse reached maximum tumor burden (*n* = 5 vehicle; *n* = 6 for 0.05 and 0.25 mg/kg ABT-700-SG3199), (**G)** Kaplan–Meier survival analysis, and **(H)** body weight measurements relative to baseline. **(I)** HCT116 CDX tumor growth until first vehicle-treated mouse reached maximum tumor burden (*n* = 4 for 0.5 mg/kg ABT-700; *n* = 5 for 0.5 mg/kg cADC; *n* = 6 for vehicle and 0.5 mg/kg ABT-700-SG3199), **(J)** Kaplan–Meier survival analysis, and **(K)** Body weight measurements relative to baseline. **(L)** Western blot and **(M)** quantification of total MET and LGR5 protein expression in DLD-1 CDX tumors at study endpoints. Statistical significance determined by log-rank test or ANOVA (*\*p* < 0.05; \**\*p* < 0.01; *\*\*\*p* < 0.001; *\*\*\*\*p* < 0.0001). Quantitative data are presented as mean ± SD.

Next, the anti-tumor activity of ABT-700-SG3199 was evaluated in DLD-1 and HCT116 cell-line derived xenograft (CDX) models. To evaluate dose dependency, DLD-1 CDXs were treated with ABT-700-SG3199 once per week for two doses at 0.05 and 0.25 mg/kg, yielding 42% and 68% TGI at day 11 when first vehicle tumor reached maximum tumor burden (**Fig. 3F**). The higher dose ABT-700-SG3199 arm showed a median survival of 43 versus 18 days for the lower dose arm (**Figs. 3F-G and Supplementary Figs. S5A-C**). Body weight remained stable across cohorts, indicating favorable tolerability (**Fig. 3H**). For HCT116 CDXs, ABT-700-SG3199 ADC, ABT-700, or cADC, were administered once per week for two doses at 0.5 mg/kg (**Figs. 3I-K and Supplementary Figs. S5D-G**). A higher dose was selected as DLD-1 tumors were not eliminated at 0.25 mg/kg (**Figs. 3F-H**). ABT-700-SG3199 showed significant TGI (83%) compared to vehicle on day 21, when the first vehicle-treated animal reached maximal tumor burden (**Fig. 3I and Supplementary Figs. S5D-G**). ABT-700 and cADC showed minimal effects. ABT-700-SG3199 also extended survival relative to vehicle, cmAb, or unconjugated ABT-700, with 2 animals surviving past 80 days (**Figs. 3J and Supplementary Figs. S5D-G**). Consistent with DLD-1 CDXs, no significant body weight changes were observed (**Fig. 3K**). These findings show ABT-700-SG3199 exhibits significant anti-tumor efficacy in MET-expressing CRC models.

### MET inhibition increases LGR5 expression in CRC cells

Since we showed that loss of LGR5 expression in response to therapy is associated with increased MET activity and/or expression in CRC cells in vitro (**Fig. 1G-I**) (10), we next examined the effect of ABT-700-SG3199 on MET and LGR5 levels. Analysis of tumors from the DLD-1 CDX study showed a dose-dependent decrease in MET and a concomitant increase in LGR5 levels following ABT-700-SG3199 treatment (**Fig. 3L-M**). Given this finding, we then tested if treatment with other MET-targeting mAbs and ADCs also increased LGR5 levels. Treatment with ABT-700 reduced total and phosphorylated MET, while simultaneously upregulating LGR5 expression in a dose-dependent manner after 24 hours in DLD-1, LoVo, and LIM1215 cells (**Fig. 4A**). Since we observed the most substantial increase in LGR5 at 0.5-1 µg/mL ABT- 700, we selected the 1 µg/mL dose to examine time-dependent effects of ABBV-399 and ABBV-400 ADCs. ABT-700-SG3199 was tested at 0.1 µg/mL due to its enhanced potency. Like ABT-700, we observed a decrease in both total and phosphorylated MET and increase in LGR5 in response to treatment with ABBV-399, ABBV-400, and ABT-700-SG3199, particularly after 48 hours (**Figs. 4B-D)**. To further interrogate whether LGR5 upregulation is attributable to MET loss or inhibition, we next examined how MET KD or MET inhibitor treatment influences LGR5 expression. DLD-1 and LoVo stable MET KD cells were generated using MET-targeting shRNA. MET KD was confirmed by western blot (**Fig. 4E**), and both DLD-1 and LoVo MET KD cells showed an increase in LGR5 expression. Similar results were observed for DLD-1 cells using MET siRNA **(Supplementary Fig. S3B)**. To assess the effect of chemical inhibition of MET on LGR5 expression, CRC cells were treated with the small molecule inhibitor Crizotinib (1 µM < IC_25_; **Supplementary Fig. S6A**). Crizotinib completely abrogated MET phosphorylation at all doses tested in a time-dependent manner (**Fig. 4F** and **Supplementary Fig. S6B**). Further, crizotinib induced a gradual increase in LGR5 levels across all three cell lines. These findings demonstrate that inhibition of MET activity, rather than changes in total MET expression, may be driving the increase in LGR5 levels. These results suggest LGR5 upregulation to be a potential mechanism of resistance to MET-targeting therapies in CRC models.

**Figure 4.**
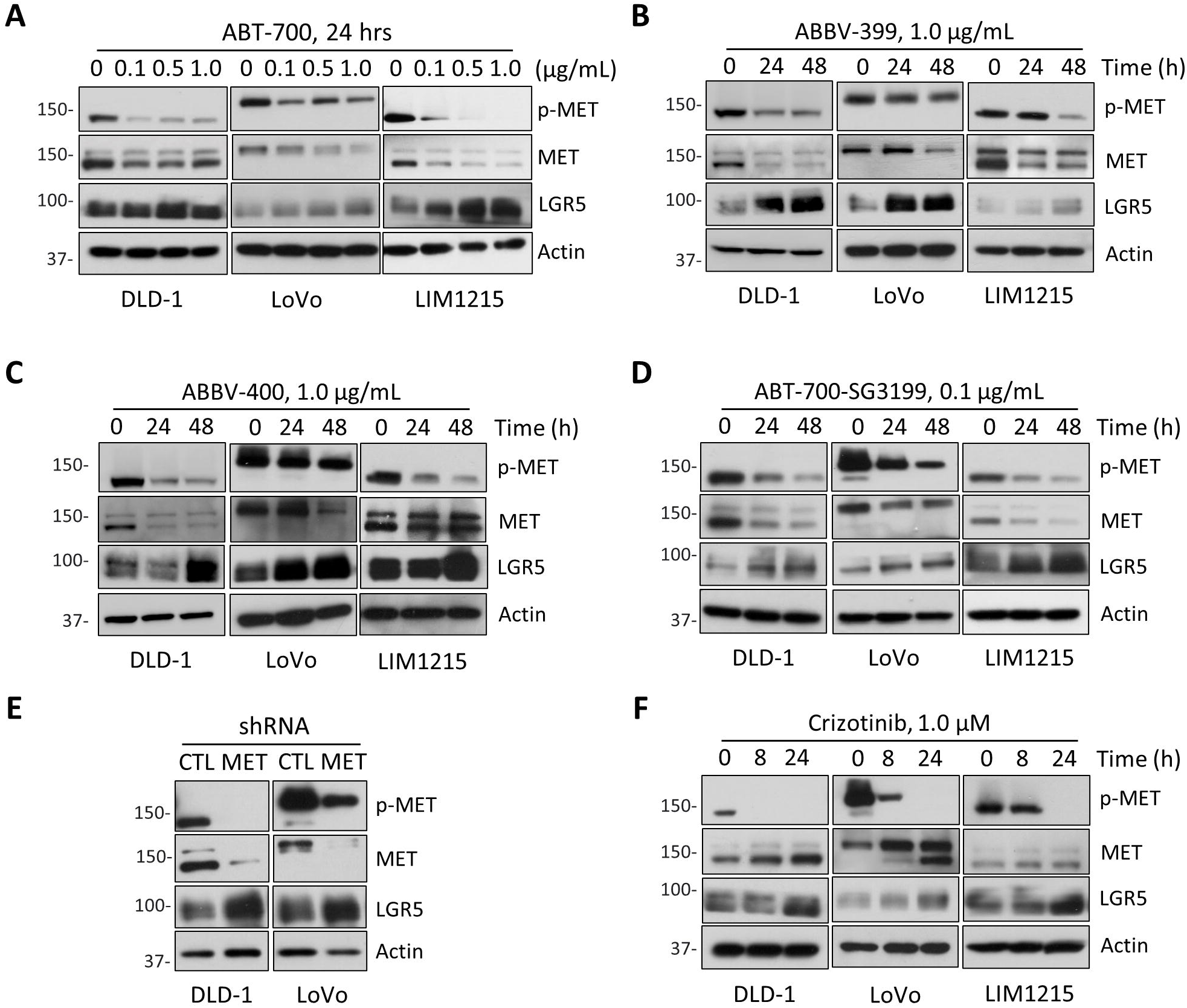
Treatment-induced loss or gene ablation of MET increases LGR5 expression in CRC cells. (A-F) Western blot analysis of LGR5, MET, and phospho-MET expression in DLD-1, LoVo, or LIM1215 cells following **(A)** dose-dependent ABT-700 treatment over 24 hours and treatment with MET-targeting ADCs **(B)** telisotuzumab vedotin (ABBV-399, 1 µg/mL), **(C)** telisotuzumab adizutecan (ABBV- 400, 1 µg/mL) and **(D)** ABT-700–SG3199 (0.1 µg/mL). **(E)** shRNA-mediated MET knockdown, and **(F)** crizotinib treatment. All treatments in **B-F** were assessed over the indicated time course. Results represent ≥3 independent experiments. Quantitative data are presented as mean ± SD.

### Dual-targeting MET and LGR5 with ADCs enhances CRC cell-killing efficacy

As our findings have shown, MET and LGR5 are inversely regulated in response to LGR5- or MET-targeting ADC treatment (**Figs. 1I, 3L-M,** and **4B-D**) (10), this suggests that dual-targeting MET and LGR5 may be a more effective strategy than either monotherapy in treating CRC. First, we tested whether MET- and LGR5 mAbs can simultaneously target and internalize into the same CRC cells. ICC and confocal microscopy demonstrated that MET and LGR5 mAbs bind and co-internalize (white arrows) in DLD-1 and LoVo cells (**Fig. 5A**) after two hours. This finding implies that by targeting both MET and LGR5 with ADCs, more payload could potentially be delivered to kill CRC cells more effectively. To evaluate the therapeutic potential of co-targeting MET and LGR5, we assessed the efficacy of combining ABT- 700-SG3199 and 8E11-CPT2 in vitro. Drug interaction analysis using the Loewe model revealed predominantly additive cytotoxicity across DLD-1, LoVo, and LIM1215 cells, with synergy emerging for the lower ABT-700-SG3199 concentrations tested in all three CRC cell lines (**Figs. 5B-D**). For example, the selected combination of 0.04 μg/mL ABT-700-SG3199 with 16.67 μg/mL 8E11-CPT2 reduced DLD-1 cell viability by 56%, compared with 30% for each monotherapy (**Fig. 5E**). In LoVo cells, combination of 0.002 μg/mL ABT-700-SG3199 with 1.85 μg/mL 8E11- CPT2 treatment yielded 54% decrease in viability versus 36% and 29% for ABT-700- SG3199 and 8E11-CPT2 monotherapy, respectively (**Fig. 5F**). In LIM1215 cells, the combination of 0.02 μg/mL ABT-700-SG3199 with 1.85 μg/mL 8E11-CPT2 resulted in 55% reduction in viability, exceeding the effects of either agent alone (**Fig. 5G**). We next evaluated whether the enhanced efficacy of the LGR5/MET ADC combination was driven by the delivery of diverse payloads (i.e., DNA-damaging SG3199 and TOP1i CPT2) or the targeting of two different antigens (i.e., MET and LGR5). First, LGR5- or MET-targeting ADC combinations, incorporating different payloads, but directed against the same target were evaluated. The efficacy of 8E11- CPT2 was tested in combination with 8E11-SG3199 (DAR=2), an LGR5-targeting ADC incorporating the same linker-payload as ABT-700-SG3199. ABT-700-SG3199 was tested in combination with ABBV-399, incorporating MMAE payload. For the latter MET combination, ABBV-399 was selected as it was the only other MET ADC that showed cytotoxicity in an LGR5-expressing CRC cell line (**Fig. 2E**). Neither the MET-nor LGR5 single-target ADC combinations showed enhanced efficacy compared to the LGR5/MET ADC combination (**Supplementary Figs. S7A-B)**. We then tested the combination of ABT-700-SG3199 and 8F2-SG3199 ADCs directed against MET and LGR5, respectively, yet incorporating the same linker-payload. However, the combination did not enhance efficacy (**Supplementary Fig. S7C**). Corresponding Loewe synergy scores are provided in **Tables S2-S3.** Together, these findings demonstrate that dual targeting of MET and LGR5 with ADCs incorporating payloads with different mechanisms of action enhances cytotoxic efficacy in vitro.

**Figure 5.**
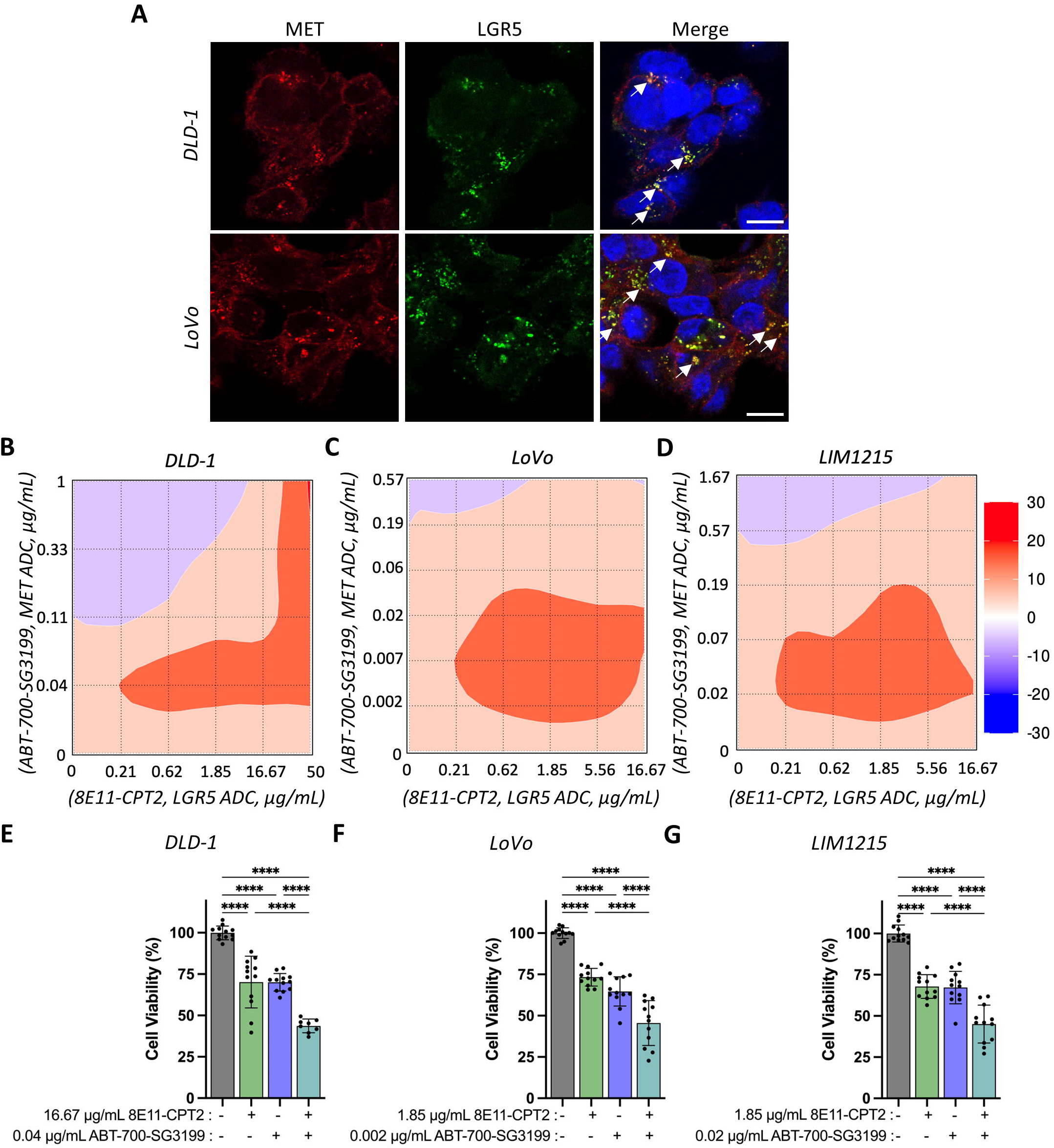
MET and LGR5 co-localization and enhanced activity of ABT-700- SG3199 in combination with 8E11-CPT2 ADC in CRC cells. **(A)** Immunocytochemistry showing internalization of ABT-700 and co-localization (white arrows) with the LGR5-targeting antibody 8F2 after 1 hour at 37 °C in MET- expressing DLD-1 and LoVo cells. Scale bars: 10 µm. Loewe synergy heat maps for **(B)** DLD-1, **(C)** LoVo, and **(D)** LIM1215 cells co-treated with increasing concentrations of ABT-700-SG3199 and 8E11-CPT2 for 5 days. Cell viability graphed for select drug combinations demonstrating additive or synergistic effects in **(E)** DLD-1, **(F)** LoVo, and **(G)** LIM1215 cells. Synergy was assessed using SynergyFinder+. Statistical significance determined by ANOVA (\*\*\**p*< 0.0001). Quantitative data are presented as mean ± SD.

### Combination treatment of MET- and LGR5-targeting ADCs exerts potent efficacy and extends survival in CRC xenograft models

To evaluate the LGR5/MET ADC combination treatment in vivo, we tested the efficacy of ABT-700-SG3199, 8E11-CPT2, or combination in MET- and LGR5-overexpressing DLD-1 CDXs (*KRAS*^MUT^), CRC-001 (*NRAS^MUT^*), and XST-GI-010 (*KRAS*^MUT^) patient-derived xenograft (PDX) models (**Figs. 6A-I and Supplementary Figs. S8A-M**) (13,24), as *RAS*^MUT^ CRC have limited treatment options compared to RAS^WT^ patients. The 8E11-CPT2 ADC, which binds both mouse and human LGR5, was shown to be well-tolerated in mice up to 20 mg/kg (17). As 0.25 mg/kg ABT-700-SG3199 significantly slowed tumor growth in DLD-1 CDXs and 0.50 mg/kg resulted in tumor stasis or regression in a fraction of HCT116 CDXs, we initially selected a slightly lower dose of 0.10 mg/kg ABT-700-SG3199 for in vivo studies to better observe combinatorial effects with 8E11-CPT2. For 8E11- CPT2, a 5 mg/kg dose was previously shown to significantly inhibit tumor growth in CRC-001 and XST-GI-010 PDX models (13). Across all three models, efficacy curves for each treatment group (Vehicle, 8E11-CPT2, ABT-700-SG3199, and ADC combo) were truncated when the first mouse reached maximal tumor burden. DLD-1 CDX models were administered a single dose of vehicle, ABT-700-SG3199 (0.1 mg/kg), 8E11-CPT2 (5 mg/kg), or the combination. The combination demonstrated improved anti-tumor efficacy and survival benefit compared to ADC monotherapy (**Figs. 6A-C and Supplementary Figs. S8B-E**). CRC-001 PDX models were administered vehicle, ABT-700-SG3199 (0.1 mg/kg), 8E11-CPT2 (5 mg/kg), or the combination once per week for two doses (**Figs. 6D-F and Supplementary Figs. S8F-I**). ABT-700-SG3199 in combination with 8E11-CPT2 showed enhanced efficacy compared to all other treatment arms, with a modest difference in TGI compared to ABT-700-SG3199 monotherapy (**Figs. 6D-F and Supplementary Figs. S8F-I**). Furthermore, the combination of ABT-700-SG3199 and 8E11-CPT2 conferred the greatest overall survival benefit (**Fig. 6E**). The XST-GI-010 PDX model was treated similarly to CRC-001, though a higher ABT-700-SG3199 dose of 0.25 mg/kg was selected to enhance anti-tumor efficacy (**Figs. 6G-I and Supplementary Figs. S8J-M**). Both ABT-700-SG3199 monotherapy and combination with 8E11-CPT2 resulted in significant TGI compared to vehicle and 8E11-CPT2 monotherapy (**Fig. 6G**). Further, the combination showed more durable response compared to monotherapy (**Figs. 6G-H**). Three animals in the combination-treated cohort-maintained stasis past day 90 (**Figs. 6G-H and Supplementary Figs. S8J- M**). No significant decrease in body weight was observed for any treatment groups across all models (**Figs. 6C, E, and I**). These findings demonstrate that ABT-700-SG3199 combined with 8E11-CPT2 results in enhanced anti-tumor efficacy and an extended survival benefit compared to either ADC monotherapy in *RAS*^MUT^ CRC models.

**Figure 6.**
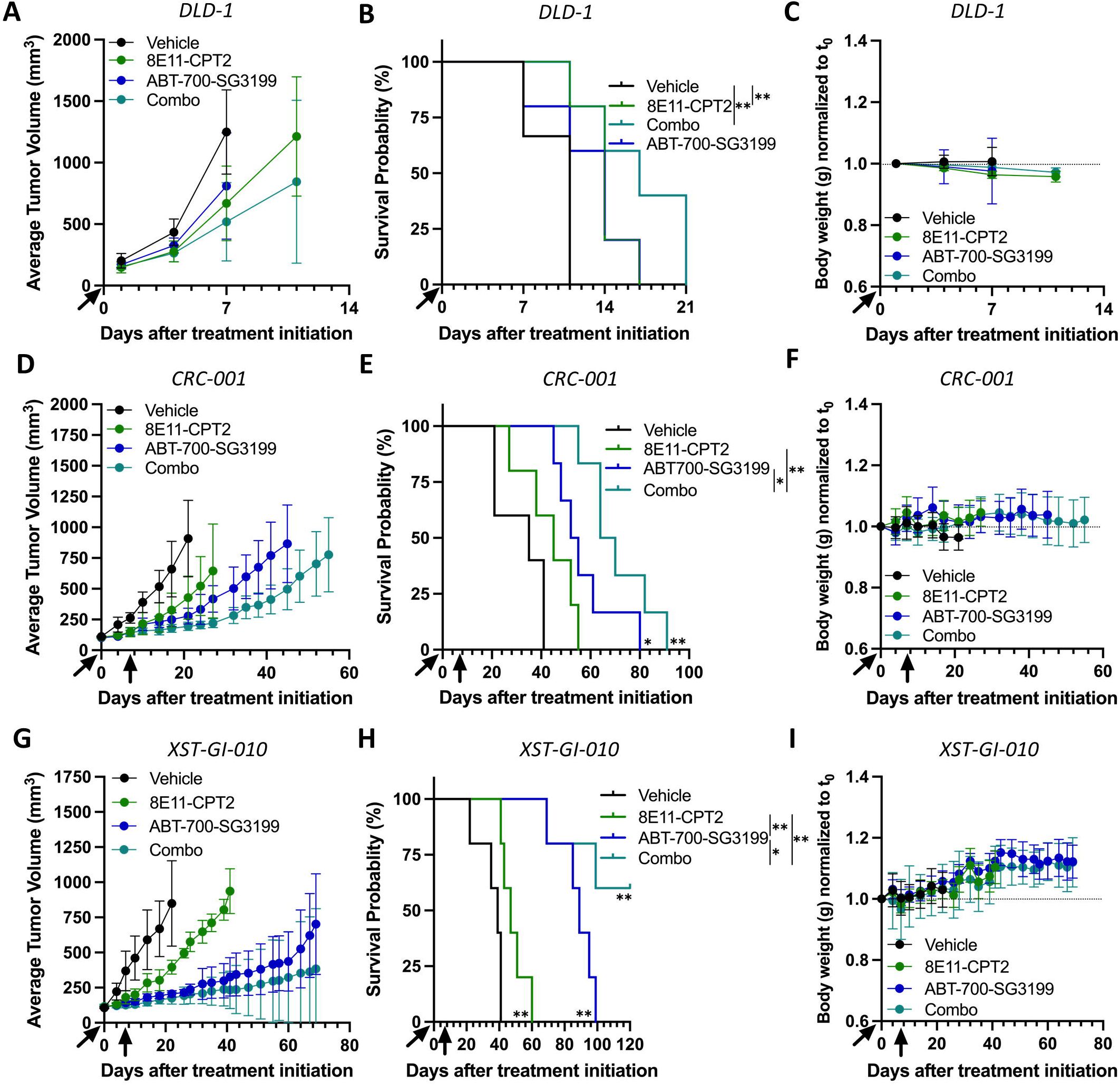
Anti-tumor efficacy and survival benefit of ABT-700-SG3199 in combination with 8E11-CPT2 in pre-clinical CRC models. DLD-1 CDX (A) tumor growth effects, (B) Kaplan–Meier survival analysis, and (C) body weight measurements relative to baseline (*n* = 6 for vehicle and *n* = 5 for all other groups). CRC-001 patient-derived xenograft (PDX) (D) tumor growth effects, (E) Kaplan–Meier survival analysis, and **(F)** body weight measurements relative to baseline (*n* = 5 for vehicle and 8E11-CPT2 and *n* = 6 for other groups). XST-GI-010 PDX **(G)** tumor growth effects, **(H)** Kaplan–Meier survival analysis, and **(I)** body weight measurements relative to baseline (*n* = 6 for vehicle and *n* = 5 for all other groups). Tumor burden measurements are cut off after the first mouse in each group reached maximum tumor burden. ADCs were administered as follows: 8E11-CPT2 at 5 mg/kg in all models and ABT-700-SG3199 at 0.1 mg/kg in DLD-1 and CRC-001 models and 0.25 mg/kg in the XST-GI-010 model. Statistical significance determined by logrank test or ANOVA ( \**p* < 0.05, \*\**p* < 0.01, \*\*\**p* < 0.001, and \*\*\*\**p* < 0.0001). Quantitative data are presented as mean ± SD.

## Discussion

Despite therapeutic advancements, CRC remains the second leading cause of cancer-related mortalities in the United States (2). Nearly 25% of CRC diagnoses present as metastatic CRC (mCRC), characterized by a 5-year survival rate of merely 15% (2). Standard-of-care chemotherapy regimens such as FOLFOX (folinic acid, 5-FU, and oxaliplatin) and FOLFIRI (folinic acid, 5-FU, and irinotecan) are hampered by low response rates, systemic toxicity, and the evolution of adaptive resistance (32). The benefits of immune checkpoint blockade are often limited to patients with Deficient Mismatch Repair or Microsatellite Instability-High (dMMR/MSI-H) tumors, a subset that accounts for only 3-5% of mCRCs (33). While targeted therapies, such as those directed against epidermal growth factor receptor (EGFR), can benefit select patient populations, their overall effectiveness can be limited by oncogenic driver mutations and both primary and acquired mechanisms of resistance (18,32). ADCs, on the other hand, have the potential to circumvent several of these drawbacks by selectively delivering cytotoxic payloads directly to tumor cells, thereby conferring potent efficacy across diverse CRC molecular subtypes irrespective of mutational status (1). In this study, we show that both chemotherapy and the LGR5-targeting ADC 8E11-CPT2 increase MET expression and activation in CRC. While our MET-targeting ADC, ABT-700-SG3199, exhibits potent activity *in vitro* and *in vivo*, recurrent tumors have increased LGR5 expression. Combining ABT-700-SG3199 with 8E11-CPT2 enhanced therapeutic potency and prolonged survival in CRC models, likely by overcoming tumor heterogeneity and resistance mechanisms.

LGR5 has emerged as a particularly compelling ADC target in CRC owing to its overexpression, established role as a marker of CSCs, and its capacity to rapidly and constitutively internalize independent of its R-Spondin 1-4 ligands (6,7,11–13,34). We confirmed LGR5 upregulation in primary and metastatic CRC tumors relative to matched normal tissue **(Figs. 1A-B)**, and significant overexpression across CRC cell lines harboring oncogenic *KRAS* and *PIK3CA* mutations **(Figs. 1D- F)**, underscoring its promise as a therapeutic target. Under cytotoxic pressure from standard-of-care agents such as 5-FU and SN-38, LGR5 expression was reduced and accompanied by upregulation and activation of MET **(Figs. 1G-H)**, extending upon previous findings by our group and others (8,10). Consistently, MET has been shown to sustain resistance to cytotoxic and targeted therapies (18,35). Building on our prior findings with MMAE-conjugated LGR5-targeting ADCs (10), treatment with 8E11-CPT2 ADC induced LGR5 downregulation and a concomitant increase in MET expression and activation in a time-dependent manner **(Fig. 1I)**, suggesting RTK-mediated activation and/or amplification as a potential resistance-bypass mechanism. Given that LGR5 KD increases chemoresistance and activates both MET and β-catenin in in CRC cells (10,36,37), and that MET is regulated by β- catenin/TCF-mediated transcriptional activation in CRC (38), this suggests a potential interplay between MET and Wnt/β-catenin signaling in coordinating CRC cell survival. Still, the exact mechanisms underlying therapy-driven loss of LGR5 warrant further investigation.

MET dysregulation is a critical driver of CRC pathogenesis (16,17), with MET overexpression detected across cell lines, primary tumors, and liver metastases **(Figs. 1A, C-E)**. However, no MET-targeting therapy has been approved for CRC patients. Numerous MET tyrosine kinase inhibitors (TKIs) have been developed to exploit MET dependency. Yet, preclinical success has not translated to clinical approval in CRC due to multi-targeting profiles and off-target toxicities (35). Anti-MET mAbs have similarly demonstrated limited efficacy due to activation of MAPK signaling or other RTK-pathways and tumor heterogeneity (23). In addition to promoting tumor progression, MET has also emerged as a driver of both primary and acquired resistance to therapies. In *RAS*^WT^ CRC, prolonged therapeutic pressure selects for MET amplification and autocrine signaling, counteracting cetuximab-induced EGFR blockade (18). MET has been shown to promote resistance to HER2- targeting ADCs and KRAS inhibitors used for the treatment of mCRC (39,40). Similarly, our results suggest that MET likely mediates resistance to LGR5-targeting ADCs, as well as to chemotherapies **(Figs. 1G-I)** (10). Given that MET upregulation may occur via transcriptional activation, post-translational modification, or both; the precise mechanism for MET upregulation and activation in response to ADC therapy remains to be further resolved. Overall, MET remains a promising therapeutic target in CRC due to its intrinsic overexpression and capacity to drive resistance to a broad spectrum of therapies.

MET-targeting ADCs improve upon prior therapeutic strategies by relying on MET overexpression to deliver cytotoxic payloads into tumor cells, independent of MET activation, mutation, or downstream signaling status (41). Validation of this approach has been supported by FDA approval of ABBV-399 (Teliso-V) in chemotherapy-refractory, MET-overexpressing non-squamous non-small cell lung cancer (NSCLC) (20,42). However, our data shows ABBV-399 exhibits limited potency in CRC cells **(Figs. 2D-G)**, as its payload MMAE is a P-glycoprotein/MDR-1 substrate subject to ABC transporter-mediated efflux, a resistance mechanism frequently encountered in CRC (37). ABBV-400 (Temeb-A), which retains the ABT-700 backbone but carries the TOP1i adizutecan, demonstrated similar activity in CRC cells **(Figs 2D-G)**. Leveraging its high affinity, target selectivity, and efficient lysosomal trafficking, we conjugated ABT-700 to an alternative, more potent payload, SG3199, to generate ABT-700-SG3199 **(Figs. 2A-C; Supplementary Figs. S2A-B and S3A)**. SG3199, or PBD, is a DNA crosslinking agent utilized in loncastuximab tesirine, a CD19-targeting ADC approved for the treatment of relapsed large B-cell lymphoma (28,31,43). ABT- 700-SG3199 exhibited potent, MET-specific cytotoxicity across CRC cell lines independent of mutation status, whereas ABBV-399 and ABBV-400 showed lower potency restricted to LoVo and HCT116 cells **(Figs. 2D-I and Supplementary Figs. S3E-G)**. Together, these results suggest that MET-targeting ADCs can be highly effective in CRC; however, ADC performance is dependent on the payload.

Although enhanced potency can improve anti-tumor activity, it may also narrow the therapeutic index (14). Thus, it is important to perform safety/toxicity studies for any new therapeutic modality. ABT-700-SG3199 was well tolerated up to the highest dose tested (1.5 mg/kg) in immunocompetent mice with no off-target payload-mediated toxicities **(Figs. 3A-E and Supplementary Figs. S4A-B)**, supporting the selection of doses between 0.05 and 0.5 mg/kg for anti-tumor efficacy evaluation in CRC CDX models. Notably, most ADC toxicities have been shown to be payload-mediated rather than target-dependent (44). The dosing range was also comparable to approved loncastuximab tesirine and pivekimab sunirine and other ADCs incorporating the same SG3199 payload (31,45). ABT-700-SG3199 induced dose-dependent tumor regression and significantly improved overall survival **(Figs. 3F-J and Supplementary Figs. S5A-H)**. In summary, these findings warrant further development of ABT-700-SG3199 as a therapeutic strategy in patient subsets with intrinsic or acquired MET overexpression, including CRC.

While ABT-700-SG3199 demonstrated durable tumor inhibition and, in some cases, regression in CRC CDX models, complete eradication as a single agent was not achieved at the tested doses **(Figs. 3G, J, and K)**. Biomarker analysis of relapsed DLD-1 CDX tumors revealed that while ABT-700-SG3199 reduced MET expression and downstream activation, LGR5 levels increased in a dose-dependent manner **(Fig. 3L-M)**. Reciprocal LGR5 upregulation was recapitulated *in vitro* following pharmacological and genetic MET abrogation in a time- and dose-dependent manner **(Figs. 4A-F and Supplementary Fig. S6B)**. Collectively, these findings suggest that MET-directed therapies either eliminate MET^high^ CRC cells with residual LGR5^⁺^/MET^low^ cells driving tumor regrowth, or LGR5 is upregulated in response to MET-targeting therapy as a mechanism of resistance, or both. In fact, induction of LGR5 expression has been shown to occur in response to cetuximab, KRAS inhibitors, and other MAPK pathway targeting agents (7,13,46–48). Collectively, these observations suggest that therapeutically targeting LGR5 could potentiate the efficacy of EGFR-, KRAS-, and MET-targeting therapies.

Combining ADCs with other systemic therapies offers a promising path forward, enabling lower dosing of each agent and more effectively targeting tumor heterogeneity and resistance. For instance, in MET-overexpressing, EGFR-mutant NSCLC, combining ABBV-399 with the EGFR TKI osimertinib produced a higher objective response rate than ADC monotherapy, offering a potential strategy to overcome MET-mediated resistance following progression on standard EGFR TKIs (49). Our group showed that pairing 8E11-CPT2 with cetuximab, which augments LGR5 levels, enhanced therapeutic potency and prolonged survival in CRC PDX models (13). Another group showed that a different PBD-based MET-targeting ADC co-administered with gemcitabine produced synergistic activity and achieved tumor regression in a pan*c*reatic cancer model (50). Our findings show that dual-targeting MET and LGR5 by combining ABT-700-SG3199 with 8E11-CPT2 produced synergistic or additive activity in CRC cell lines **(Figs. 5B-G)**. The ADC combination enhanced tumor inhibition and prolonged survival relative to either single-agent ADC **(Figs. 6A-I and Supplementary Figs. S8B-M)**, suggesting that simultaneous engagement of two distinct antigens and delivery of complementary payloads improves antitumor efficacy by ensuring effective payload delivery even when LGR5 or MET expression is variable. Furthermore, neither MET-targeted ADCs with different payloads nor MET- and LGR5-targeted ADCs sharing the same PBD payload produced robust cancer cell killing in vitro **(Supplementary Figs. S7A-C).** The latter observation is consistent with emerging clinical data on sequential ADC dosing, which suggests that target switching offers limited additional benefit when the ADCs share a similar payload (51). Collectively, our results suggest that MET- and LGR5-directed ADC combinations have the potential to overcome both tumor heterogeneity and adaptive resistance.

While our findings establish proof of principle for MET- and LGR5-directed ADC combination therapy in CRC, several considerations warrant further investigation. The bidirectional MET–LGR5 crosstalk described here requires deeper mechanistic dissection. As ABT-700-SG3199 does not cross-react with murine MET, we were unable to evaluate on-target toxicity, though doses tested were well below those used clinically for other PBD-based ADCs (31). Comprehensive pharmacokinetic, pharmacodynamic, and safety evaluation in non-human primates will therefore be essential to define the therapeutic window of the ABT-700-SG3199 + 8E11-CPT2 combination. Additionally, testing this dual-targeting strategy using different doses in orthotopic models of mCRC spanning diverse mutational backgrounds and molecular subtypes will be necessary. Future work should also explore the ideal ADC dosing concentrations, frequency, and sequential rather than concurrent administration to establish the optimal therapeutic window.

In summary, we have identified a bidirectional MET–LGR5 signaling axis that may drive ADC resistance in CRC, validating that dual-targeting these receptors may serve as a rational therapeutic strategy to improve CRC treatment. We generated ABT-700-SG3199, a MET-targeting PBD-based ADC, and comprehensively evaluated it as both a monotherapy and in combination with the LGR5-targeting ADC, 8E11-CPT2, where the combination demonstrated enhanced anti-tumor activity and extended survival in preclinical CRC models. Overall, these findings establish preclinical proof of concept for MET- and LGR5-directed ADC combinations and lay the foundation for next-generation dual-targeting strategies, including METxLGR5 bispecific antibodies and ADCs, in CRC and other MET- and LGR5- expressing malignancies.

## Data availability

Publicly available RNA sequencing (RNA-seq) datasets analyzed in this study were obtained from The Cancer Genome Atlas (TCGA) colorectal adenocarcinoma (COADREAD) cohort via cBioPortal (http://www.cbioportal.org) and from the Gene Expression Omnibus (GEO) under accession number GSE50760. All raw data generated during this study are available at Mendeley Data (doi: 10.17632/6wyrkrg74x.1) and from the corresponding author upon request.

## Authors’ Contributions

Conceptualization: K.S.C.; Methodology: K.S.C., S.S., P.C.H.; Investigation: S.S., P.C.H., Z.L., C.G.-B., M.G.C., S.P.S., K.S.C.; Data curation: S.S., P.C.H., Z.L., C.G.-B., M.G.C., S.P.S., A.M.A., L. L., S.P., K.S.C.; Formal Analysis: S.S., P.C.H., C.G.-B., L.L., S.P., K.S.C.; Visualization: S.S., P.C.H., K.S.C., Validation: S.S. and K.S.C., Writing-original draft: S.S.; Writing-review and editing: S.S., P.C.H., C.G.-B., M.G.C., S.P.S., K.S.C; Resources: K.S.C.; Funding acquisition: K.S.C.; Supervision: K.S.C.

## Acknowledgements

This work was supported by funding from NIH/NCI (R21 CA302991, R21 CA282378, R01 CA226894, and R01 CA281962) and the Jerold B. Katz Endowment in Stem Cell Research to K.S.C., a predoctoral fellowship in the Biomedical Informatics, Genomics, and Translational Cancer Research Training Program (BIG-TCR) funded by CPRIT (RP210045) to S.S, a predoctoral fellowship of the Gulf Coast Consortia, on the Training Interdisciplinary Pharmacology Scientists Program (T32 GM139801) and Andrew Sowell-Wade Huggins Fellowship from The University of Texas MD Anderson Cancer Center UTHealth Houston Graduate School of Biomedical Sciences to P.C. H., and a predoctoral Schissler Foundation Fellowship from The University of Texas MD Anderson Cancer Center UTHealth Houston Graduate School of Biomedical Sciences to C.G-B. We would like to thank Dr. Julie Rowe, Betty Arceneaux, and Dr. Karan Saluja with assistance in patient sample collection and processing, and Martha Thompson for assistance with regulatory approvals for research involving human subjects. Schematics were created with Biorender.com.

