## Supplementary Tables S1-S3 for "Antibody-drug conjugate combination therapy targeting LGR5 and MET with different payloads enhances efficacy in preclinical colorectal cancer models"

| **ABT-700-SG3199, IC_50_** | | |
| --- | --- | --- |
| **Cell line** | **ng/ml** | **pM** |
| DLD-1 | 7.18 ± 2.78 | 48.0 ± 18.6 |
| LoVo | 1.20 ± 0.64 | 8.03 ± 4.29 |
| LIM1215 | 5.67 ± 3.16 | 37.9 ± 21.1 |
| HCT116 | 2.10 ± 0.83 | 14.0 ± 5.57 |
| SW620 | 2.11 ± 0.41 | 14.1 ± 2.72 |
| LS180 | 12.0 ± 5.72 | 80.4 ± 38.2 |
| HT-29 | 1.31 ± 1.19 | 8.77 ± 7.99 |

**Supplementary Table S1. Average IC_50_ values for ABT-700-SG3199 in a panel of colorectal cancer cell lines. Related to Figure** **2.** Results representative of ≥3 independent experiments performed in triplicate. Quantitative data are presented as mean ± SD.

| **LOEWE SYNERGY SCORES** | | | | | | | |
| --- | --- | --- | --- | --- | --- | --- | --- |
| **Treatments** | | **DLD-1** | | | | | |
| **8E11-CPT2, μg/mL** | | **0.21** | **0.62** | **1.85** | **5.56** | **16.67** | **50** |
| **ABT-700- SG3199, μg/mL** | **1** | NT | -8.31 | -4.69 | -1.97 | 2.19 | 21.14 |
|  | **0.33** |  | -4.84 | -2.13 | 0.56 | 6.7 | 18.95 |
|  | **0.11** |  | -1.57 | 1.89 | 7.27 | 7.44 | 16.04 |
|  | **0.04** |  | 10.59 | 14.05 | 12.91 | 12.79 | 13.68 |
| **Treatments** | | **LoVo** | | | | | |
| **8E11-CPT2, μg/mL** | | **0.21** | **0.62** | **1.85** | **5.56** | **16.67** | **50** |
| **ABT-700-**  **SG3199, μg/mL** | **0.57** | -1.24 | -0.85 | 0.57 | 1.19 | -0.4 | NT |
|  | **0.19** | 0.67 | 1.97 | 3.66 | 2.35 | 3.36 |  |
|  | **0.06** | 4.89 | 7.82 | 8.27 | 6.84 | 5.14 |  |
|  | **0.02** | 6.09 | 12.39 | 11.74 | 11.12 | 10.86 |  |
|  | **0.007** | 10.47 | 15.65 | 17.45 | 17.29 | 9.44 |  |
|  | **0.002** | 7.31 | 11.87 | 15.57 | 13.55 | 6.67 |  |
| **Treatments** | | **LIM1215** | | | | | |
| **8E11-CPT2, μg/mL** | | **0.21** | **0.62** | **1.85** | **5.56** | **16.67** | **50** |
| **ABT-700-**  **SG3199, μg/mL** | **1.67** | -8.26 | -7.25 | -4.47 | -1.47 | 2.29 | NT |
|  | **0.56** | -1.81 | 1.2 | 4.41 | 5.5 | 4.73 |  |
|  | **0.19** | 5.17 | 5.35 | 9.58 | 9.6 | 7.05 |  |
|  | **0.06** | 10.22 | 10.25 | 13.45 | 12.22 | 6.69 |  |
|  | **0.02** | 12.04 | 14.5 | 18.83 | 13.26 | 9.9 |  |

**Supplementary Table S2. Loewe synergy scores for ABT-700-SG3199 in combination with 8E11-CPT2 ADC. Related to Figure 5.** NT, not tested. Results representative of the average of 3 independent experiments performed in quadruplicate.

| **LOEWE SYNERGY SCORES** | | | | | | |
| --- | --- | --- | --- | --- | --- | --- |
| **Treatments** | | **LIM1215** | | | | |
| **8E11-SG3199, μg/mL** | | **1** | **0.5** | **0.1** | **0.05** | **0.01** |
| **8E11-CPT2, μg/mL** | **20** | -3.84 | -2.3 | 1.1 | NT | NT |
|  | **5** | -5.91 | -5.74 | 5.59 |  |  |
|  | **1** | -2.76 | -2.9 | 4.75 |  |  |
| **Treatments** | | **LoVo** | | | | |
| **ABT-700-SG3199, μg/mL** | | **0.556** | **0.185** | **0.062** | **0.021** | **0.007** |
| **ABBV-399, μg/mL** | **20** | -38.74 | -45.5 | -42.04 | -39.34 | -41.64 |
|  | **6.7** | -39.99 | -57.82 | -49.18 | -59.16 | -46.79 |
|  | **2.2** | -34.57 | -50.68 | -58.25 | -49.36 | -38.4 |
|  | **0.74** | -17.48 | -26.86 | -35.42 | -36.61 | -39.72 |
| **Treatments** | | **LoVo** | | | | |
| **ABT-700-SG3199, μg/mL** | | **0.556** | **0.185** | **0.062** | **0.021** | **0.021** |
| **8F2-SG3199, μg/mL** | **0.556** | 0.56 | 1.51 | 1.37 | 1.37 | 2.02 |
|  | **0.185** | -1.27 | -1.16 | 0.17 | 0.14 | 0.42 |
|  | **0.062** | -8.83 | -4.54 | -3.76 | -0.98 | -0.85 |
|  | **0.021** | -17.93 | -19.45 | -15.75 | -1.88 | -2.88 |
|  | **0.007** | -24.2 | -27 | -13.95 | -2.69 | -1.43 |
|  | **0.002** | -33.79 | -50.1 | -42.24 | -27.91 | -13.95 |
|  | **0.0008** | -37.87 | -50.65 | -48.73 | -25.11 | -11.42 |

**Supplementary Table S3. Loewe synergy scores for MET and LGR5 ADC combinations with same or different payloads. Related to Supplementary Figure S7.** NT, not tested.
