## Supplementary Files S1-S8 for "Antibody-drug conjugate combination therapy targeting LGR5 and MET with different payloads enhances efficacy in preclinical colorectal cancer models"

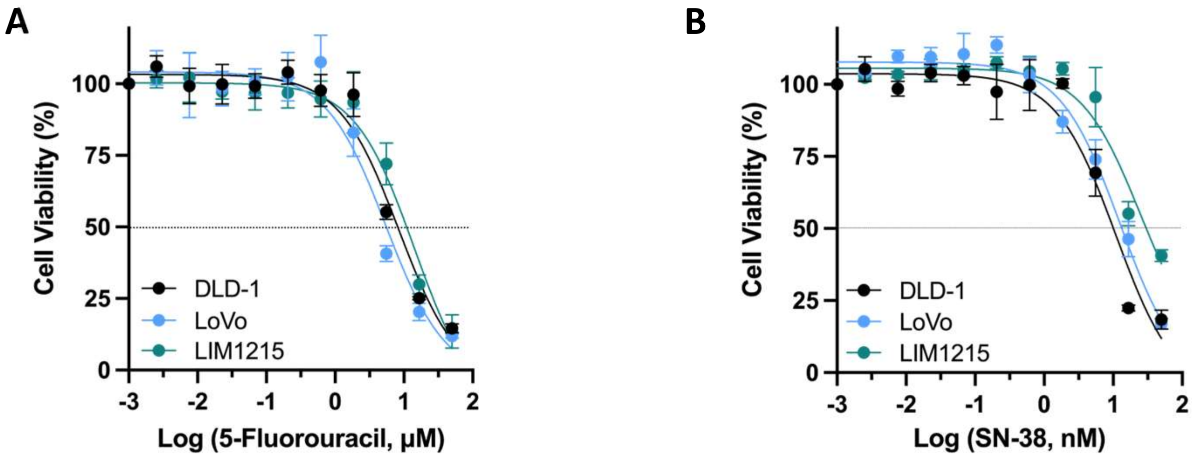


**Supplementary Figure S1. Cytotoxicity of standard-of-care chemotherapies in CRC cell lines. Related to Figure 1.** Sensitivity of DLD-1, LoVo, and LIM1215 cells to **(A)** 5-fluorouracil (5-FU) and **(B)** SN-38. Experiments were performed 2–3 times in triplicate; cytotoxicity was measured 5 days post-treatment. Quantitative data are presented as mean ± SD.

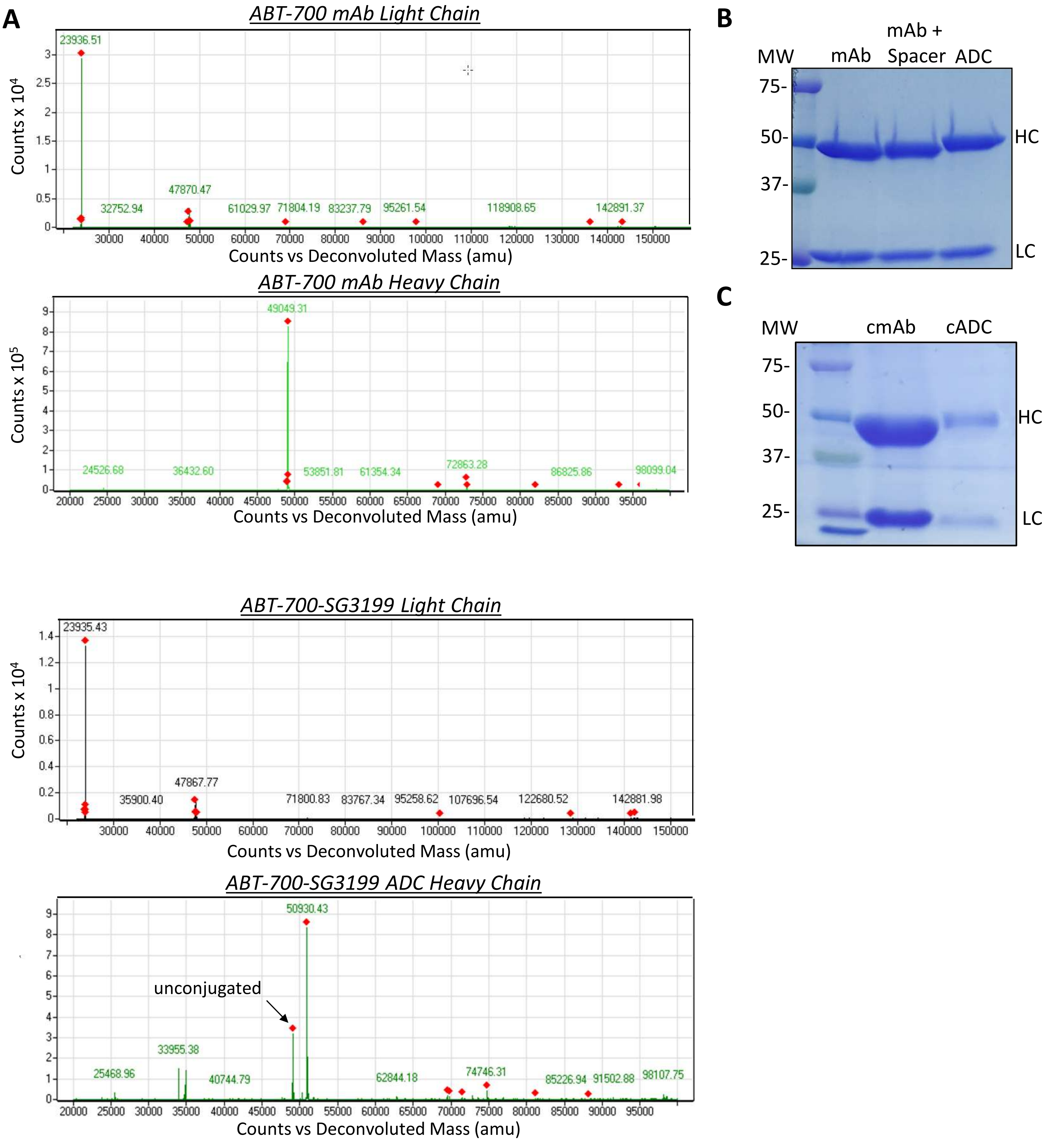


**Supplementary Figure S2. Mass spectrometric analysis and Coomassie staining of non-targeting cADC and ABT-700-SG3199 ADC. Related to Figure 2. (A)** LC-MS analysis of ABT-700 mAb and ABT-700-SG3199 ADC light and heavy chains. Coomassie staining of **(B)** ABT-700 and **(C)** control (R20) mAbs and their corresponding ADCs under reducing conditions. HC: Heavy Chain; LC: Light Chain; MW: Molecular Weight.

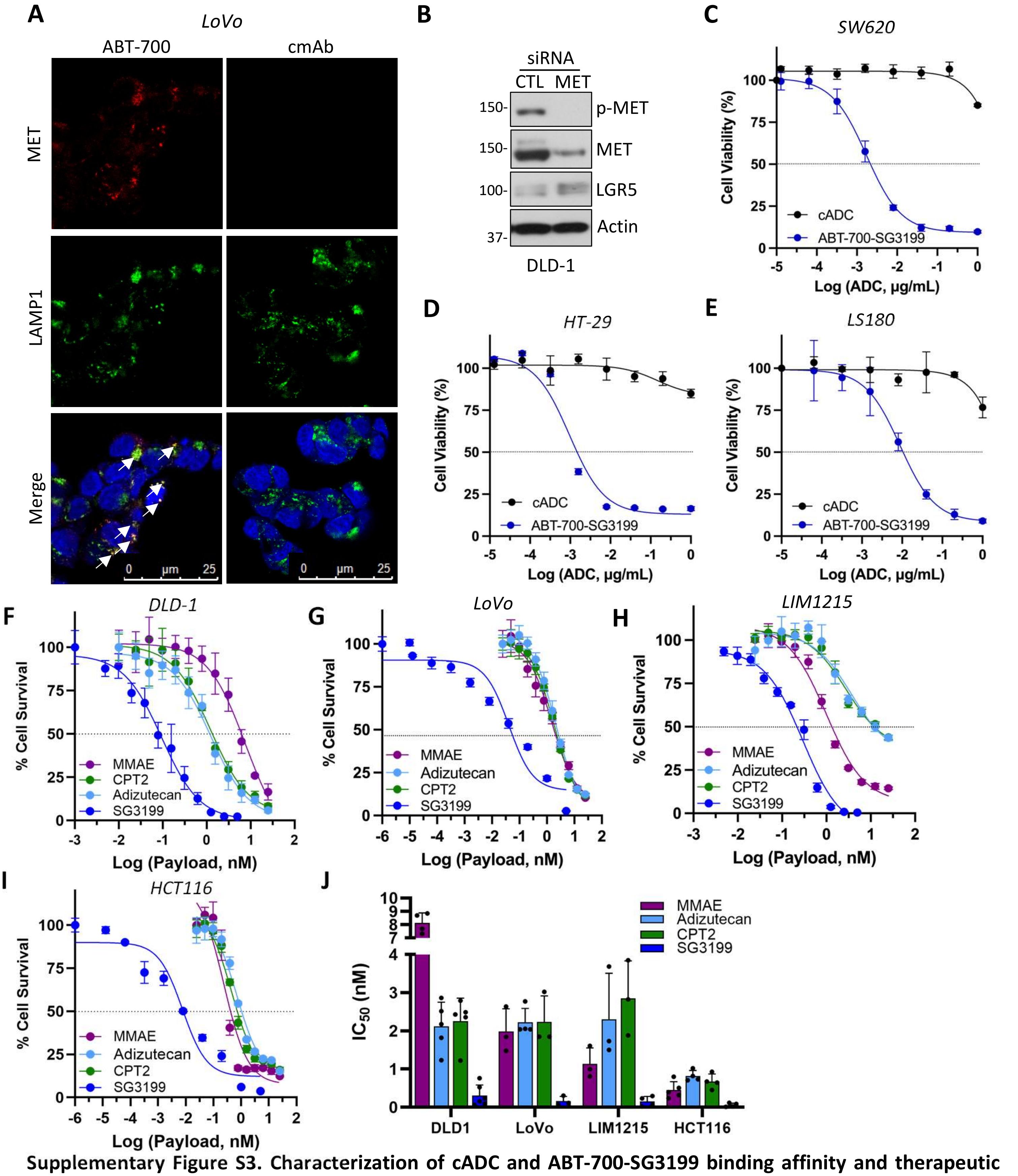


**Supplementary Figure S3. Characterization of cADC and ABT-700-SG3199 binding affinity and therapeutic potency in CRC cell lines. Related to Figure 2. (A)** Confocal microscopy confirming ABT-700 colocalization (white arrows) with LAMP1⁺ lysosomes in LoVo cells. Scale bars: 25 µm. **(B)** Western blot analysis of phospho-MET, total MET, and LGR5 protein levels in control and MET-knockdown (KD) DLD-1 cells. **(C-E)** Efficacy of ABT-700-SG3199 compared to cADC in **(C)** SW620, **(D)** HT-29 and **(E)** LS180 CRC cell lines. Results represent ≥3 independent experiments. Quantitative data are presented as mean ± SD. Sensitivity of MMAE, CPT2, Adizutecan, and SG3199 payloads after 3 days in **(F)** DLD-1, **(G)** LoVo, **(H)** LIM1215, and **(I)** HCT116 CRC cells. **(J)** IC50 values of free payloads across the CRC cell line panel. Results represent ≥3 independent experiments. Quantitative data are presented as mean ± SD.

**A**

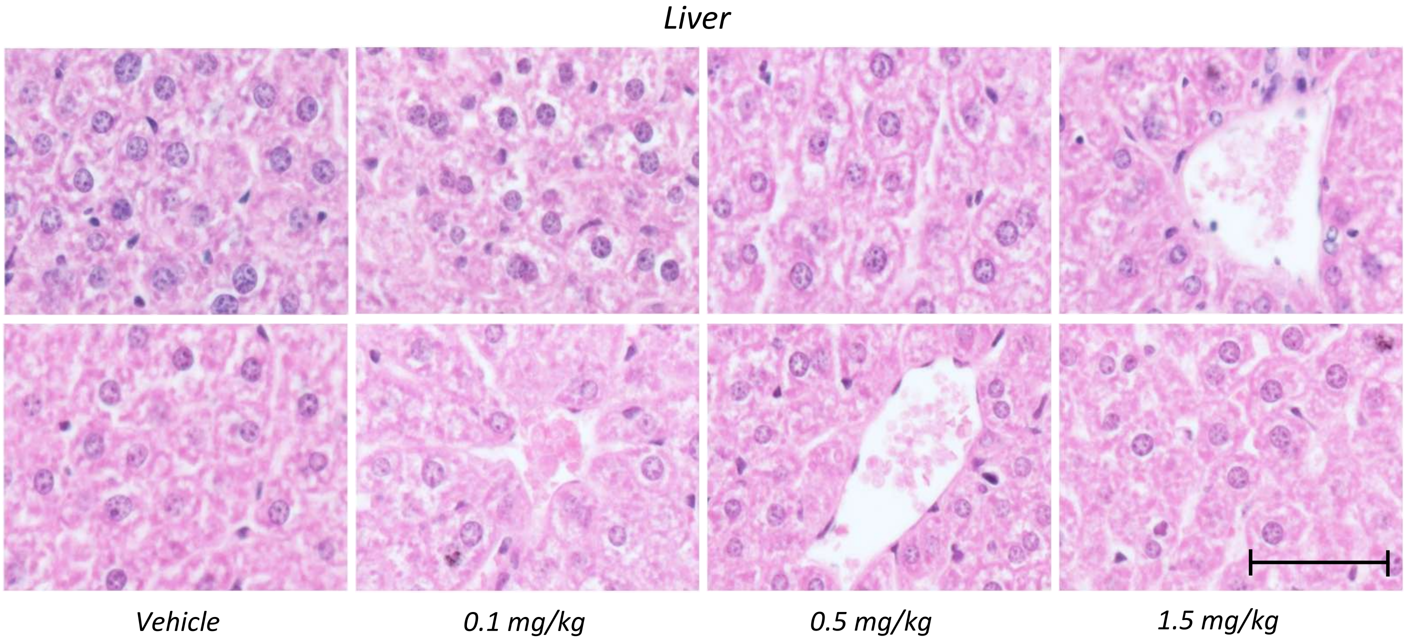


**B**

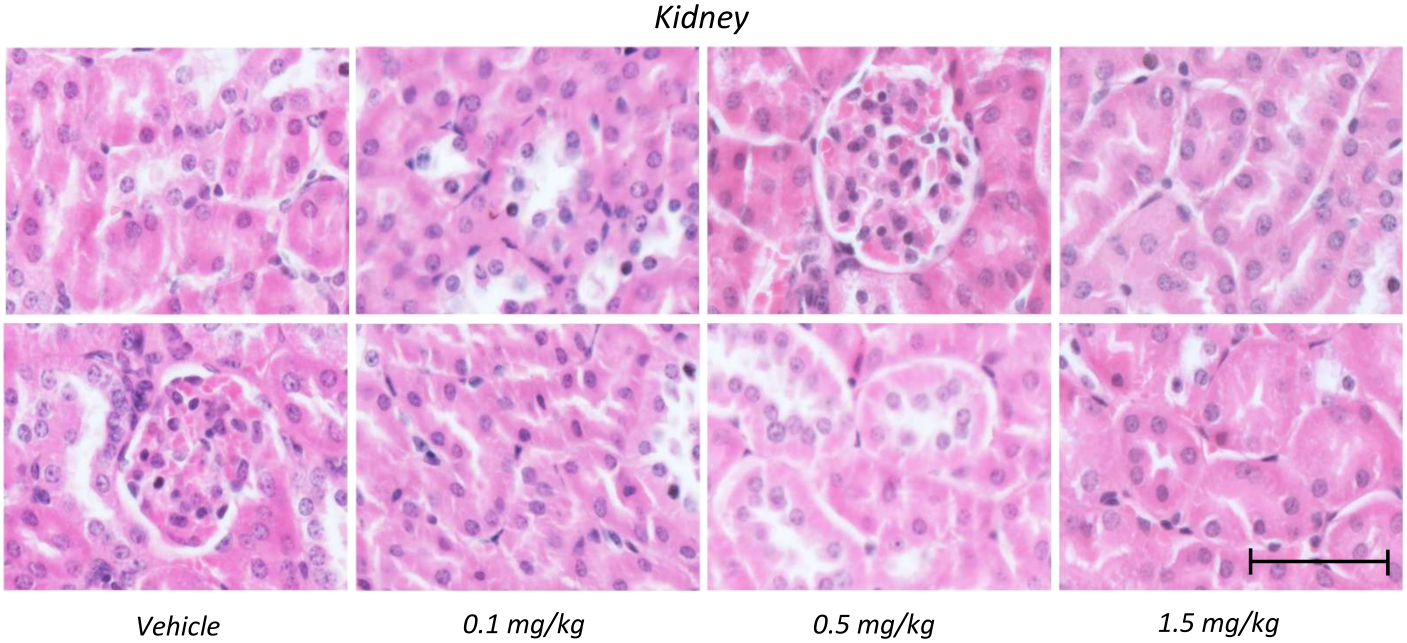


**Supplementary Figure S4. Histopathological analysis of ABT-700–SG3199 ADC tolerability in kidney and liver. Related to Figure 3. (A-B)** H&E staining of **(A)** hepatic and **(B)** renal tissues from two representative C57BL/6 mice 2 weeks after administration of ABT-700-SG3199 at the indicated doses or PBS vehicle, showing no abnormal morphological changes. Images were acquired at 20X magnification. Scale bars: 100 µm.

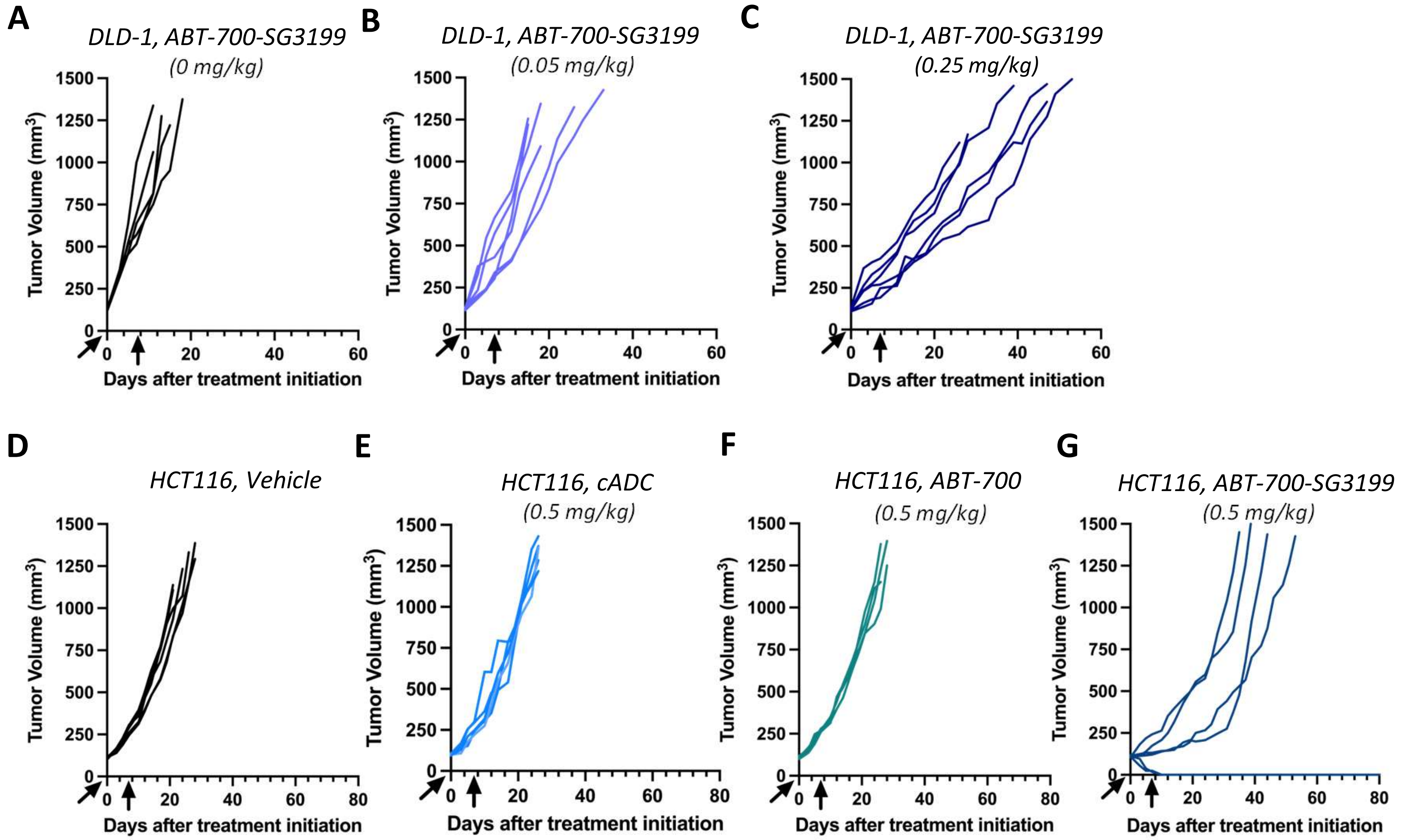


**Supplementary Figure S5. Individual tumor growth curves from CDX efficacy studies. Related to Figure 3.** Anti-tumor efficacy per mouse graphed for **(A)** Vehicle, **(B)** 0.05 mg/kg and **(C)** 0.25 mg/kg ABT-700-SG3199 cohorts dosed IP in DLD-1 CDX mice. **(D)** Vehicle, **(E)** cADC, **(F)** ABT-700 and **(G)** ABT-700–SG3199 cohorts dosed IP at 0.5 mg/kg in HCT116 CDX mice.

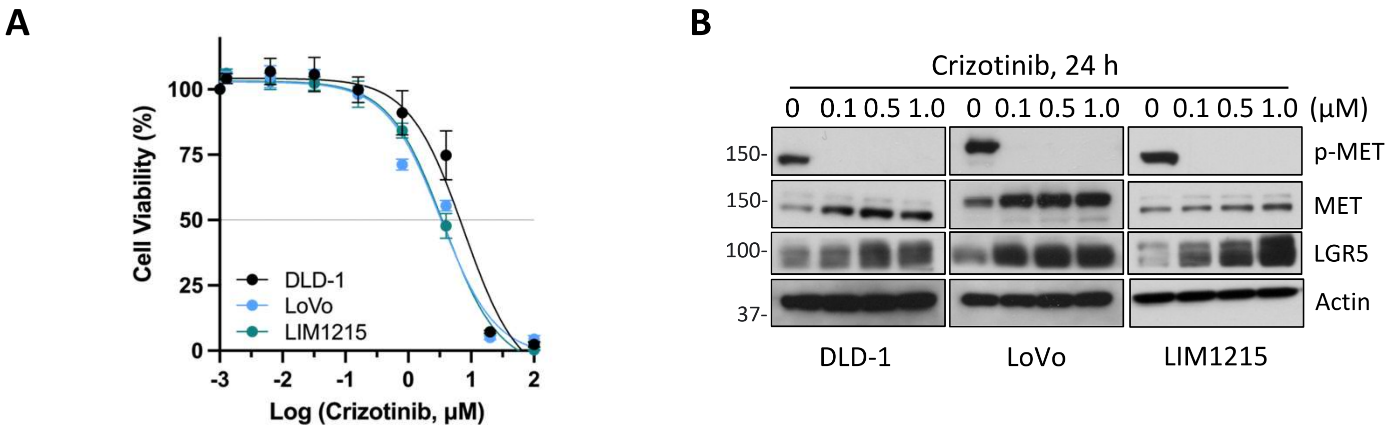


**Supplementary Figure S6. Crizotinib exhibits micromolar potency and upregulates LGR5 expression in CRC cells. Related to Figure 4. (A)** Dose-dependent sensitivity of DLD-1, LoVo, and LIM1215 cells to Crizotinib, **(B)** Western blots of changes in total MET, phospho-MET and LGR5 expression in DLD-1, LoVo, and LIM1215 cells following dose-dependent Crizotinib treatment over 24 hours.

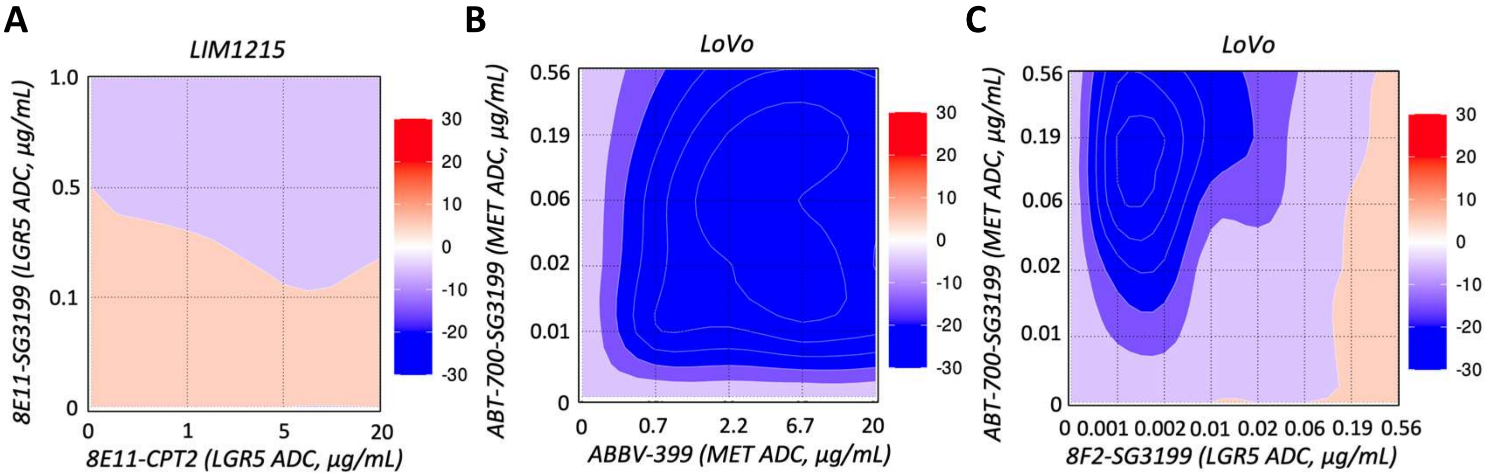


**Supplementary Figure S7. Synergism analysis of MET and LGR5 single-target ADC combinations in CRC cells. Related to Figure 5.** Loewe synergy heat maps for **(A)** LIM1215 cells co-treated with LGR5 ADCs with differing payloads, **(B)** LoVo cells co-treated with MET ADCs with varying payloads, and **(C)** LoVo cells co-treated with MET- and LGR5 ADCs incorporating different payloads for 5 days.

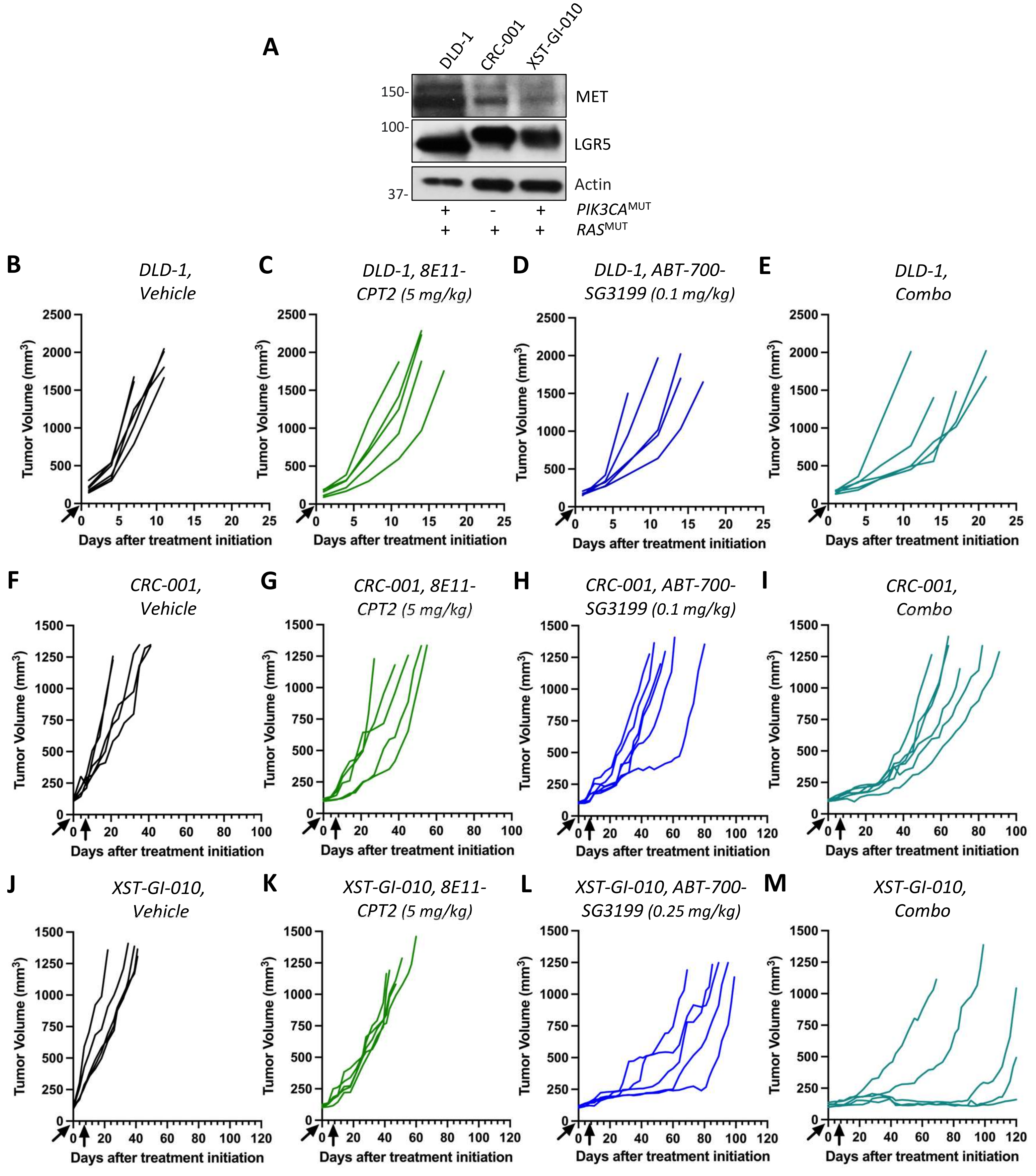


**Supplementary Figure S8. individual tumor growth curves from ABT-700-SG3199 and 8E11-CPT2 combination studies in CDX and PDX models. (A)** Western blots of endogenous MET and LGR5 in PDX and DLD-1 CDX models. All harbor *KRAS*$^{MUT}$, except *NRAS*$^{MUT}$ CRC-001. **(B-M)** Anti-tumor efficacy per mouse graphed for **(B-E)** DLD-1 CDX mice, **(F-I)** CRC-001**,** and **(J-M)** XST-GI-010 PDX models treated with vehicle, 8E11-CPT2, ABT-700-SG3199 and ADC combo. ADCs were administered IP: 8E11-CPT2 at 5 mg/kg in all mouse studies, and ABT-700-SG3199 at 0.1 mg/kg in DLD-1 and CRC-001 studies and 0.25 mg/kg in XST-GI-010 study.
